# Inhibition of tryptophan 2,3-dioxygenase impairs DNA damage tolerance and repair in glioma cells

**DOI:** 10.1101/2020.05.28.110874

**Authors:** Megan R. Reed, Leena Maddukuri, Amit Ketkar, Stephanie D. Byrum, Maroof K. Zafar, April C. L. Bostian, Alan J. Tackett, Robert L. Eoff

## Abstract

Aberrant expression of tryptophan 2,3-dioxygenase (TDO) is a determinant of malignancy and immune response in gliomas in part through kynurenine (KYN)-mediated activation of the aryl hydrocarbon receptor (AhR). In the current study, we investigated the hypothesis that TDO activation in gliomas has a broad impact upon genome maintenance - promoting tolerance of replication stress (RS) and repair of DNA damage. We report that inhibition of TDO activity attenuated recovery from hydroxyurea (HU)-induced RS and increased the genotoxic effects of bis-chloroethylnitrosourea (BCNU), as fork progress was impeded when TDO-deficient glioma cells were treated with BCNU. Activation of the Chk1 arm of the replication stress response (RSR) was reduced when TDO activity was blocked prior to treatment with BCNU, whereas phosphorylation of serine 33 (pS33) on replication protein A (RPA) was enhanced – indicative of increased fork collapse. Restoration of KYN levels protected against some replication-associated effects of BCNU. Inhibition of TDO activity had a strong anti-proliferative effect on glioma-derived cells – enhancing the cytotoxic effects of BCNU. Analysis of results obtained using quantitative proteomics revealed TDO-dependent changes in several signaling pathways – including down-regulation of DNA repair factors and sirtuin signaling. Consistent with these observations, inhibition of TDO diminished SIRT7 recruitment to chromatin, which increased histone H3K18 acetylation – a key mark involved in 53BP1 recruitment to sites of DNA damage. Cells lacking TDO activity exhibited defective recruitment of 53BP1 to gH2AX foci, which corresponded with delayed repair of BCNU-induced DNA breaks. Addition of exogenous KYN increased the rate of break repair. The discovery that TDO activity modulates sensitivity to DNA damage by fueling SIRT7/53BP1 localization to chromatin and repair of BCNU-induced DNA damage highlights the potential for tumor-specific metabolic changes to influence genome stability and may have implications for glioma biology and treatment strategies.

## INTRODUCTION

Glioblastoma multiforme (GBM or simply glioblastoma) represents an especially deadly type of primary brain tumor afflicting both adult and pediatric patients (1–3). The challenges to effective treatment of glioblastoma are many and include the location of the tumor, invasive microtubes, tumor heterogeneity, a relatively high proportion of tumor initiating (or stem-like) cells, and a robust replication stress/DNA damage response (RSR/DDR) capacity (4–7). The factors driving increased tolerance of DNA damage and replication stress (RS) are likewise multi-factorial and strongly correlated with resistance to genotoxic drugs and tumor recurrence in glioblastoma patients (8).

The aberrant and constitutive degradation of tryptophan to kynurenine (KYN) and subsequent activation of the AhR (a ligand-activated transcription factor involved in a variety of biological processes) has emerged as a driving force in multiple aspects of GBM biology (9). One route to KYN pathway (or KP) activation in GBMs occurs in response to COX2/prostaglandin E_2_-mediated up-regulation of tryptophan 2,3-deoxygenase (TDO2 or TDO) (10). TDO is one of three human enzymes catalyzing the rate-limiting step in the conversion of tryptophan to KYN. In GBM and advanced stage breast cancer, TDO promotes a pro-malignant/anti-immune response through production of KYN, an endogenous agonist of the AhR transcription factor, and other tryptophan catabolites. The KP-AhR cascade produces different effects on tumor and immune cells. To date, almost all studies on the KP in cancer have focused on immunosuppressive effects. The use of epacadostat (an inhibitor of KP signaling) in combination with Merck’s anti-PD-1 antibody pembrolizumab in phase III clinical trials for the treatment of advanced stage melanoma, highlights the interest in targeting the KP as an adjuvant to immune checkpoint inhibitors (11), although it remains uncertain if targeting KP-related enzymes is a useful therapeutic strategy (12).

Previously, we showed that modulation of the KP affected the expression of the translesion synthesis (TLS) enzyme DNA polymerase kappa (hpol κ) (13). Treating GBM-derived cell lines with an inhibitor of TDO lowered hpol κ expression and led to a diminished level of micronuclei (MN). The same was true for GBM cells treated with either the AhR antagonist CH-223191 or siRNA against hpol κ. Combined inhibition/knock-down of either TDO and AhR or TDO and hpol κ did not decrease MN levels further – suggestive of an epistatic relationship. Since hpol κ performs multiple functions related to DNA damage tolerance and the resolution of RS, and the AhR is a transcription factor known to regulate a variety of pathways, we postulated that attenuation of the KP might sensitize GBM cells to genotoxic drugs by changing the basal RSR/DDR capacity.

In the current study, we have investigated the hypothesis that RSR and DDR programs in glioma-derived cells are responsive to KP signaling and that this connection modulates sensitivity to RS and DNA damage. We tested this idea by measuring KP-dependent changes to DNA replication and repair in glioma-derived cells treated with either hydroxyurea (HU) or the DNA damaging agent bis-chloroethylnitrosourea (BCNU or Carmustine). We focused on analyzing changes related to RSR/DDR activation and fork dynamics, as well as more direct readouts for genomic integrity (e.g., alkaline comet assay, micronucleation assay). Cell cycle progression, viability, and motility were also assessed. Quantitative proteomic analysis was used to evaluate KP-related changes in an unbiased and global manner. Cumulatively, our results are consistent with the notion that activation of KYN signaling increased the threshold for tolerance of DNA damage and RS in GBM cells – a phenomenon that could have implications for genotoxic therapies.

## RESULTS

### Inhibition of TDO leads to diminished resolution of RS induced by either HU or the bis-functional DNA damaging agent BCNU

Since gliomas (especially GBM) exhibit remarkably high levels of RS and given our previous findings with hpol κ, we first examined the connection between KP signaling and replication dynamics. We investigated the effect of TDO inhibition on fork rate using the DNA fiber spreading (DFS) assay. We monitored DNA synthesis before exposure to BCNU (CldU, red tracks), as well as in the presence of BCNU (IdU, green tracks; **Fig. 1A**). A decrease in the ratio of IdU/CldU track lengths is indicative of fork slowing in response to treatment during the second pulse. Treating T98G cells with 680C91 for 24 h prior to the addition of IdU/CldU did not alter the rate of DNA synthesis (**Fig. 1B**). As expected, treatment with BCNU during the second pulse (IdU) reduced the fork rate by ~30% (**Fig. 1B**). Inhibition of TDO activity enhanced the effect of BCNU on fork progression, as evidenced by another ~15% decrease in the IdU/CldU ratio relative to treatment with BCNU alone (**Fig. 1B**). We attempted to further modify fork rate by adding exogenous KYN (60 μM) to the cells. While addition of KYN alone did not impact fork progression, it did protect against BCNU-mediated fork slowing when KYN was added prior to treatment with the DNA damaging agent (**Fig. 1B**). These results are consistent with the idea that modulating KP signaling alters the capacity of T98G cells to effectively replicate DNA in the face of BCNU-induced damage – with higher levels of KYN promoting damage bypass and TDO inhibition leading to increased fork stalling.

**Figure 1.**
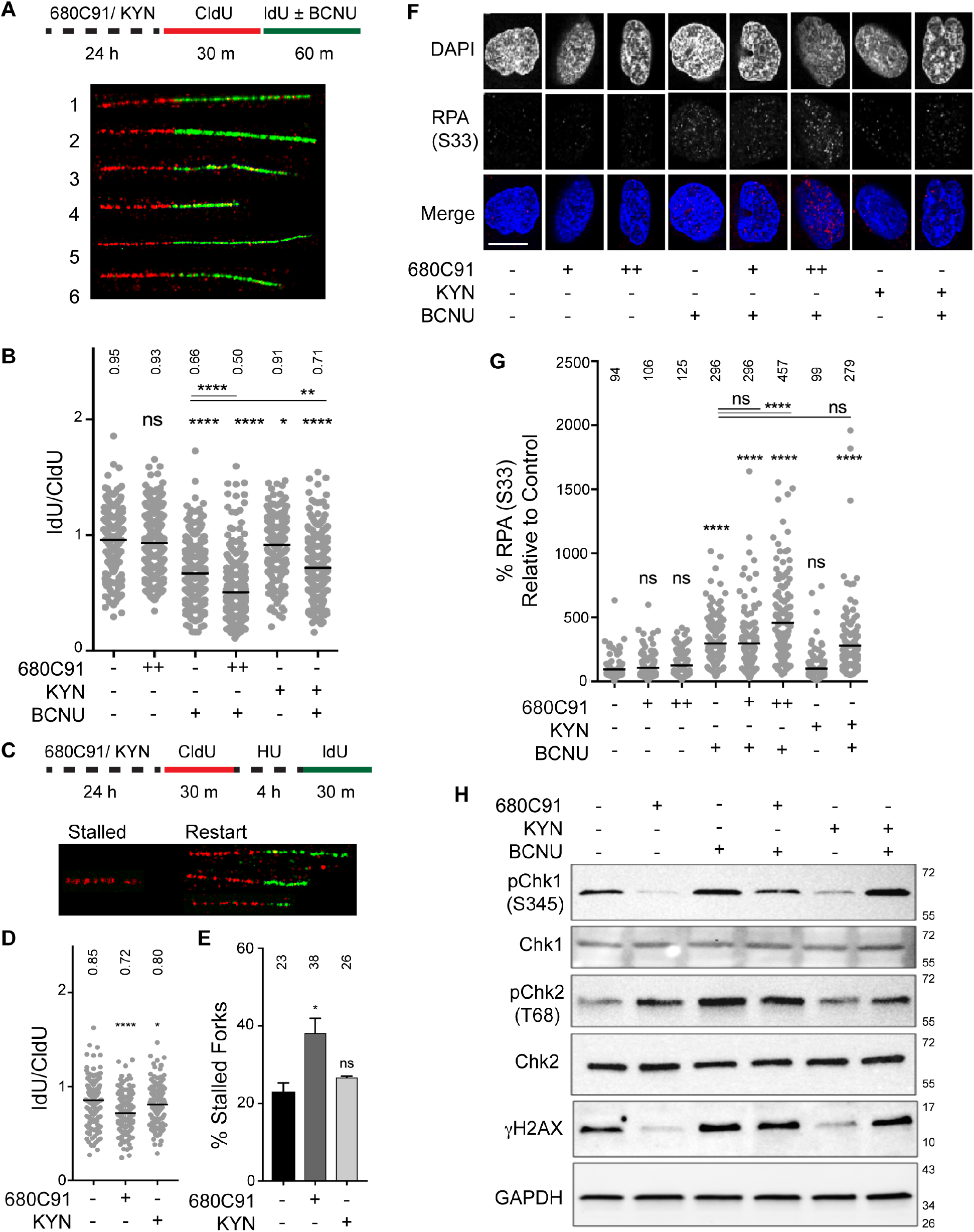
Inhibition of TDO leads to diminished resolution of stress at replication forks. *A*, Cartoon schematic of analogue treatments and representative fiber images; 1. CTL, 2. 680C91 (20 μM), 3. BCNU (125μM), 4. BCNU + 680C91, 5. KYN (60 μM) and 6. BCNU + KYN. *B*, Quantification of IdU/CldU ratio, where at least 290 fibers were scored from three biological replicates and mean ratio is depicted above each condition. CldU and IdU track lengths were normalized to individual pulse times prior to attaining IdU/CldU ratio. *C*, Cartoon schematic and representative fiber images of stalled and restarted forks. *D*, IdU/CldU ratio for restarted forks, where at least 179 fibers were scored from two biological replicates and mean ratio is depicted above each condition. For all DNA fiber spreading experiments, statistical significance was calculated using a Mann-Whitney U test, where * is P ≤ 0.05, ** is P ≤ 0.01, *** is P ≤ 0.001 and **** is P ≤ 0.0001. *E*, Percentage of stalled forks after pre-treatment with 680C91 or KYN. The mean (± s.d.) is shown with the mean percentage of stalled forks for each condition is shown above the histogram. Statistical analysis was performed using a Student’s *t* test where * is P ≤ 0.05. *F*, Representative immunofluorescence images of pS33-RPA2, in cells treated +/- 680C91, KYN, and BCNU. Scale bar: 10 μm. *G*, Quantification of pS33-RPA2 fluorescence intensity relative to the control for the above-mentioned treatments, where at least 130 cells were quantified from three biological replicates. A one-way ANOVA with a Tukey post-test was used for statistical analysis, where * is P≤ 0.05, ** is P≤ 0.01, *** is P ≤ 0.001 and **** is P ≤ 0.0001. *H*, Immunoblot of cells treated +/- 680C91/ KYN/ BCNU, and probed for; pChk1 (Ser-345), pChk2 (Thr-68), γH2AX (Ser-139) and GAPDH.

We next determined if GBM-derived cells with attenuated KP could recover from hydroxyurea (HU)-induced RS (**Fig. 1C-E**). Similar to replication defects observed with BCNU, we found that 680C91 pretreatment impaired fork restart (**Fig. 1D**) and increased the fraction of forks stalled by HU treatment (**Fig. 1E**). Pre-treating cells with KYN did not alter fork recovery from HU-induced replication stress one way or another (**Fig. 1D** and **E**). From these experiments, we concluded that T98G cells lacking TDO activity were more susceptible to HU-induced RS, consistent with the notion that KP signaling helps sustain an effective RSR in GBM-derived cells. With these results in hand, we went on to study the impact of TDO activity on markers of RSR and DDR programs.

We used immunofluorescence (IF) microscopy to monitor formation of phospho-S33 RPA2 (RPA32) in T98G cells. The serine 33 residue of RPA2 is one of the phosphatidylinositol 3-kinase related kinase (PIKK) consensus sites phosphorylated by the ATR kinase in response to replication fork stalling, facilitating stabilization of stalled forks and resolution of RS intermediates through the recruitment of factors, such as PALB2 and BRCA2, to sites of DNA damage/replication stress (14, 15). In addition to pS33 RPA2, we also monitored changes in pS345 Chk1, pT68 Chk2, and gH2AX levels via immunoblotting. In this way, we hoped to discern whether TDO activity impacts ATR signaling either through Rad17-mediated Chk1 activation, which occurs in response to fork slowing/stalling or subsequent Nbs1-mediated ATR signaling and corresponds with accumulation of pS33 RPA2 and more extensive end-resection near collapsed forks (16). We used Chk2 phosphorylation and gH2AX as indicators of DNA break formation.

Treatment of cells with the TDO inhibitor 680C91 (10 or 20 μM for 24 h) did not alter the baseline signal intensity for pS33 RPA2 in GBM-derived T98G cells (**Fig. 1F** and **G**). There was a notable decrease in pS345 Chk1 for cells treated with 680C91 (20 μM for 24 h), while pT68 Chk2 formation increased (**Fig. 1H**). Similar to the RSR marker pChk1, the DNA damage marker γH2AX decreased following treatment with 680C91 (**Fig. 1H**). The loss of pChk1 signal that accompanied TDO inhibition is interesting given that hpol κ has been shown to help activate the Rad17-arm of the ATR signaling cascade leading to phosphorylation of the Chk1 kinase. Previously we observed down-regulation of hpol κ in response to TDO inhibition (13). In this respect, the reduction in pS345 Chk1 when TDO was inhibited might be due to suppression of hpol κ/Rad17-mediated activation of ATR (16, 17).

As expected, treating cells with BCNU (125 μM, 24 h) increased the pS33 RPA2 signal, consistent with an elevation in DNA damage-induced RS (**Fig. 1G**). There was a concomitant increase in pChk1, pChk2, and γH2AX levels in response to BCNU treatment (**Fig. 1H**). Pre-treating cells with 10 μM 680C91 did not alter the level of pS33 RPA2 formed in response to BCNU treatment, but the addition of 20 μM 680C91 prior to BCNU exposure led to noticeably higher levels of pS33 RPA2 (**Fig. 1G**), which could be interpreted as a diminished capacity for resolving BCNU-induced RS in cell lacking active KP signaling. Consistent with this notion, there was noticeable suppression of pChk1 activation when cells treated with 20 μM 680C91 were subsequently exposed to BCNU (**Fig. 1H**). Activation of pChk2 and γH2AX formation following BCNU treatment was reduced slightly by TDO inhibition but not to the extent observed for pChk1 (**Fig. 1H**). It is possible that TDO inhibition suppressed RSR activation through the hpol κ-Rad17 arm of ATR/Chk1 signaling, which led to an increased reliance on pS33 RPA2 accumulation and Nbs1-mediated ATR activation in response to BCNU-induced DNA damage. This is at least consistent with the clearly higher levels of pS33 RPA2 that coincide with noticeable reduction in pChk1 activation.

Adding exogenous KYN (60 μM, 24 h) did not change basal pS33-RPA2 levels in T98G cells (**Fig. 1G**). The addition of KYN reduced pS345 Chk1 and γH2AX relative to DMSO but did not alter pT68 Chk2 levels (**Fig. 1H**). When combined with BCNU treatment, exogenous KYN seemed to at least sustain Chk1 S345 phosphorylation and γH2AX levels, but the relative level of Chk2 activation in cells pre-treated with KYN was kept below that observed for BCNU alone (**Fig. 1H**). By way of comparison with 680C91, activating the KP with exogenous KYN seemed to maintain ATR/Chk1 signaling and limit ATM/Chk2 activation following treatment with BCNU whereas blockade of TDO activity shifted the damage response away from Chk1 arm of the ATR response – leading to an accumulation of pS33 RPA2. These findings are consistent with the idea that an active KP promotes the effective resolution of fork stress in glioma-derived cells.

### BCNU-induced nuclear γH2AX signal intensity was increased by TDO inhibition

Next, we investigated whether blockade of TDO activity had an effect on the level of nuclear γH2AX in GBM-derived T98G cells treated with BCNU. To more closely examine the relationship between TDO activity and γH2AX, we treated T98G cells with either 10 (+) or 20 μM (++) 680C91 for 24 h and measured nuclear γH2AX signal intensity by IF microscopy (**Fig. 2A**). Similar to our results with whole cell lysates, there was a decrease in nuclear γH2AX signal intensity in 680C91-treated cells, but at the selected concentrations the change was modest (**Fig. 2B**).

**Figure 2.**
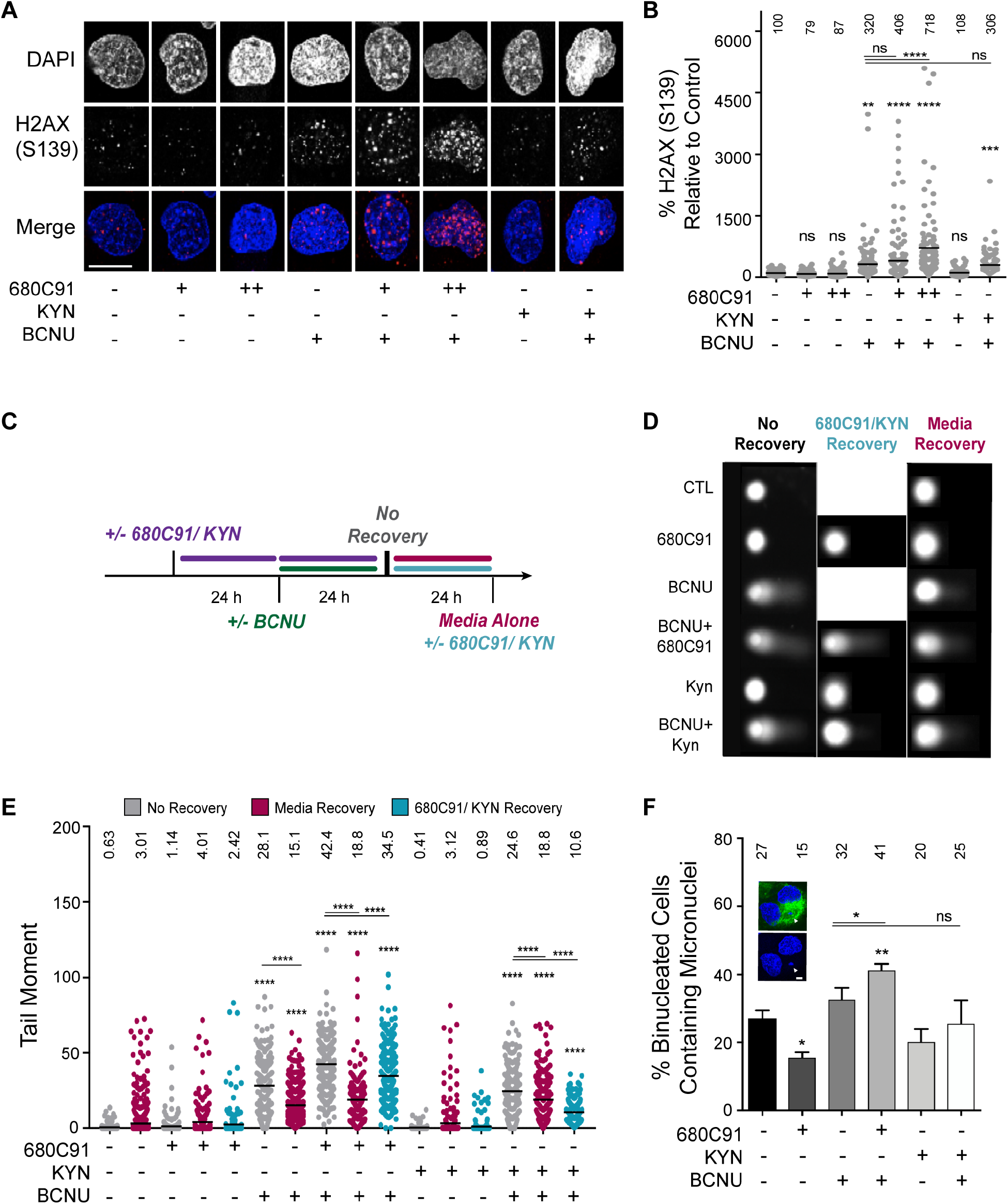
TDO inhibition results in delayed repair of BCNU-induced strand breaks and elevated chromosomal abnormalities. *A*, Representative immunofluorescence images of γH2AX (pSI39) in cells +/- 680C91, KYN and BCNU. Scale bar: 10 μm. Cells were treated with 680C91 (+, 10 μM; ++, 20 μM), KYN (60 μM), BCNU (125 μM) or a combination of these compounds. *B*, Quantification of γH2AX fluorescence intensity, where at least 160 cells were quantified from three biological replicates. *C*, Cartoon of experimental scheme for monitoring DNA strand break formation with the alkaline comet assay. *D*, Representative comet images of no recovery conditions or conditions in which treated cells were allowed to recover from BCNU in 680C91/KYN or in media alone. Cells were treated with 680C91 (20 μM), KYN (60 μM), BCNU (125 μM) or a combination of these compounds. The mean normalized intensity value for each condition is shown above the data points. *E*, Tail moment for experiments outlined in panels *C* and *D*, where at least two biological replicates were performed and at least 170 cells were scored. The mean tail moment value is shown above the data points. *F*, Representative image of a binucleated cell containing micronuclei, and quantification of the percentage of binucleated cells containing micronuclei. Cells were pre-treated with 680C91 (20 μM) or KYN (60 μM) for 24 h prior to treatment with BCNU (125 μM) or media for another 24 h. The mean percentage of binucleated cells with MN is shown above the histogram for each experimental condition. At least three biological replicates were performed. Scale bar: 5 μm. For all panels, statistical analysis was calculated using a one-way ANOVA with a Tukey post-test, where * is P≤ 0.05, ** is P≤ 0.01, *** is P ≤ 0.001 and **** is P ≤ 0.0001.

We next tested for BCNU-induced changes in nuclear γH2AX. As expected, exposing cells to BCNU (125 μM, 24 h) increased the nuclear γH2AX signal detected by IF (**Fig. 2B**). The combined effect of TDO inhibition and BCNU treatment led to a marked and dose-dependent increase in gH2AX signal intensity by IF microscopy (**Fig. 2B**). The increase in gH2AX signal observed by IF was not apparent from immunoblots with whole cell lysates. However, subsequent experiments support the notion that inhibition of TDO resulted in an accumulation of gH2AX on DNA following exposure to BCNU.

Adding KYN back to the cells did not have a major impact on nuclear gH2AX signal intensity (**Fig. 2B**). When exogenous KYN was added prior to treatment with BCNU, there was a slight (~5%) reduction in gH2AX detected by IF relative to BCNU alone (**Fig. 2B**). In summary, the changes in gH2AX measured using IF microscopy were largely in-line with the 680C91-dependent alteration in pS33 RPA2 signal observed in response to BCNU treatment (**Fig. 1G**), suggestive of an increase in unresolved DNA damage when TDO-deficient cells were exposed to BCNU.

### TDO inhibition slowed the rate of break repair following exposure to BCNU but the addition of exogenous KYN enhanced break repair

Given the effects observed for replication dynamics and DDR activation, we next used the alkaline comet assay to determine if modulation of KP signaling led to altered levels of strand breaks (**Fig. 2C** and **D**). We allowed the cells to grow in the presence of DMSO (CTL), 680C91 (20 μM), or KYN (60 μM) for 24 h before adding BCNU (or DMSO) to the media for an additional 24 h and then measuring strand breakage (**Fig. 2C**, “no recovery”). In the absence of DNA damage, culturing cells in either 680C91 or KYN alone did not have a significant impact on DNA strand breaks relative to the control, although 680C91 did increase the mean tail moment from 0.63 to 1.14 and adding KYN decreased the tail moment slightly to 0.41 (**Fig. 2E**, results presented as gray data points).

Exposing cells to BCNU (125 μM, 24 h) increased the tail moment to 28.1, more than 40-fold over that observed for DMSO treated T98G cells (**Fig. 2E**, gray data points for + BCNU). Strikingly, pretreatment with 680C91 increased the strand breaks induced by BCNU to 42.4, another 1.5-fold above that measured for BCNU alone (**Fig. 2E**, gray data points for + 680C91 + BCNU). Adding exogenous KYN to the cells prior to treatment with BCNU had a slight protective effect, as evidenced by a tail moment of 24.6 (**Fig. 2E**, gray data points for + 680C91 + KYN), which was less than the tail moment of 28.1 observed for BCNU alone.

The initial comet assay results led us to wonder if continued modulation of TDO activity and KP signaling would impact DNA repair once BCNU was removed. To investigate this possibility, we measured strand-break formation in cells treated as before except that we allowed an additional 24 h recovery period after BCNU was removed. Repair was allowed to proceed in either media (DMSO/CTL), 680C91 (20 μM), or KYN (60 μM). In this way, we were able to determine if KP signaling impacted the kinetics of break repair.

First, we controlled for repair of endogenous strand-breaks by measuring the tail moment from cells grown for 48 h in the presence of DMSO (CTL), 680C91, or KYN and allowed to recover an additional 24 h in media. Allowing the DMSO-treated cells to grow an additional 24 hours in increased the tail moment from 0.63 to 3.01 (**Fig. 2E**, compare gray control in the first column with the magenta data points in the second column). Culturing T98G cells for 48 h in the presence of 680C91 followed by a 24 recovery in media alone increased the tail moment slightly compared to the DMSO-treated control (**Fig. 2E**, compare a tail moment of 3.01 for the untreated control with a tail moment of 4.01 for + 680C91). Cells grown for 48 h in the presence of exogenous KYN followed by a 24 recovery in media alone did not alter the tail moment relative to the DMSO-treated control (**Fig. 2E**, compare a tail moment of 3.01 for the untreated control with a tail moment of 3.12 for + KYN).

For cells treated with BCNU alone, the additional recovery period reduced the tail moment ~45%, from 28.2 to 15.1 (**Fig. 2E**, comparing BCNU alone in gray with BCNU alone + 24 h media recovery in magenta), indicative of active repair of BCNU-induced strand-breaks. For cells pre-treated with the TDO inhibitor prior to BCNU, removing 680C91 and allowing cells to recover in media alone reduced the tail moment ~55%, from 42.5 to 18.8 (**Fig. 2E**, compare gray to magenta for + BCNU + 680C91). Culturing GBM cells with exogenous KYN and BCNU, followed by recovery in media reduced the tail moment from 24.6 to 18.8 (**Fig. 2E**, compare gray data points with magenta data points for + BCNU + KYN). In short, cells treated with BCNU and allowed to recover in media for 24 h exhibited a reduction in the number of strand breaks and this reduction was most pronounced for cells that had been exposed to the TDO inhibitor – perhaps indicative of a scenario where restoration in TDO activity stimulated DNA repair.

Next, we allowed the cells to recover in the presence of either 680C91 or KYN. Allowing the cells to grow for an additional 24 h in the presence of 680C91 alone (i.e., no BCNU) did not change the tail moment much compared to recovery in media (**Fig. 2E**, compare the tail moment of 4.01 for recovery in media to a tail moment of 2.42 for 72 h culture in the presence of 680C91, light blue data points). However, continuous inhibition of TDO during the recovery period prevented the efficient repair of BCNU-induced strand breaks, as evidenced by the fact that the tail moment decreased less than 20% from 42.5 to 34.6 (**Fig. 2E**, compare gray to light blue for + BCNU + 680C91). This was compared to the tail moment of 18.8 observed for cells exposed to BCNU and 680C91 then allowed to recover in media.

The effects on break repair for cells allowed to recover an additional 24 h in exogenous KYN were also interesting. Recall that there was a reduction in tail moment from 24.6 to 18.8 or ~25% when cells exposed to exogenous KYN and BCNU were then allowed to recover in media (**Fig. 2E**, right side of plot – compare gray to magenta for + BCNU + KYN). When cells were allowed to recover in the presence of exogenous KYN, there was an even larger decrease in the tail moment (24.6 to 10.6 or ~55%; far-right side of **Fig. 2E –** compare gray to light blue for + BCNU + KYN). These results support the idea that excess KYN increased the rate of repair of BCNU-induced DNA strand breaks. This is in contrast to the delayed repair of DNA breaks observed when TDO activity was blocked.

### BCNU-induced chromosomal damage was enhanced in cells with attenuated KP activity

To monitor the effect of KP signaling on chromosomal instability (CIN), we analyzed changes in MN. Treating T98G cells with 680C91 (20 μM, 24 h) reduced the number of binucleated cells with MN from 27% to 15% (**Fig. 2F**), similar to what we reported previously with a lower dose of 680C91. Treating cells with BCNU increased the percentage of cells with MN to 33% as expected (**Fig. 2F**). The percentage of binucleated cells with MN increased to 41% when 680C91 treatment preceded exposure to BCNU (**Fig. 2F**), in line with the comet assay results. The addition of exogenous KYN (60 μM, 24 h) reduced MN formation slightly compared to the DMSO control, although the *P*-value was >0.05 (**Fig. 2F**). Similarly, adding KYN resulted in a slight (but non-significant) protection from BCNU-induced MN (**Fig. 2F**, compare BCNU alone to BCNU + KYN). In summary, modulating KP signaling through inhibition of TDO had a notable impact on CIN in GBM-derived cells but the addition of exogenous KYN produced very little if any change in the formation of BCNU-induced CIN.

### KYN signaling alters progression through the S and G2/M checkpoints in BCNU-treated cells

We next sought to determine if combining KP modulation with DNA damage had an influence on cell cycle progression. Treating T98G cells with 10 μM 680C91 had minimal effect on cell cycle distribution, but treatment with 20 μM 680C91 seemed to exert a modest anti-proliferative effect on T98G cells (**Fig. 3A, 3B** and **S1**). At 20 μM 680C91, there was a slight decrease in the fraction of cells in S-phase (>2N) – from 29.3% for the untreated control to 25.4% for 680C91 treated cells - and a slight increase in the sub-G1 (<2N) population - from 0.6% for untreated cells to 1.7% for cells exposed to 20 μM 680C91 (**Fig. 3B**).

**Figure 3.**
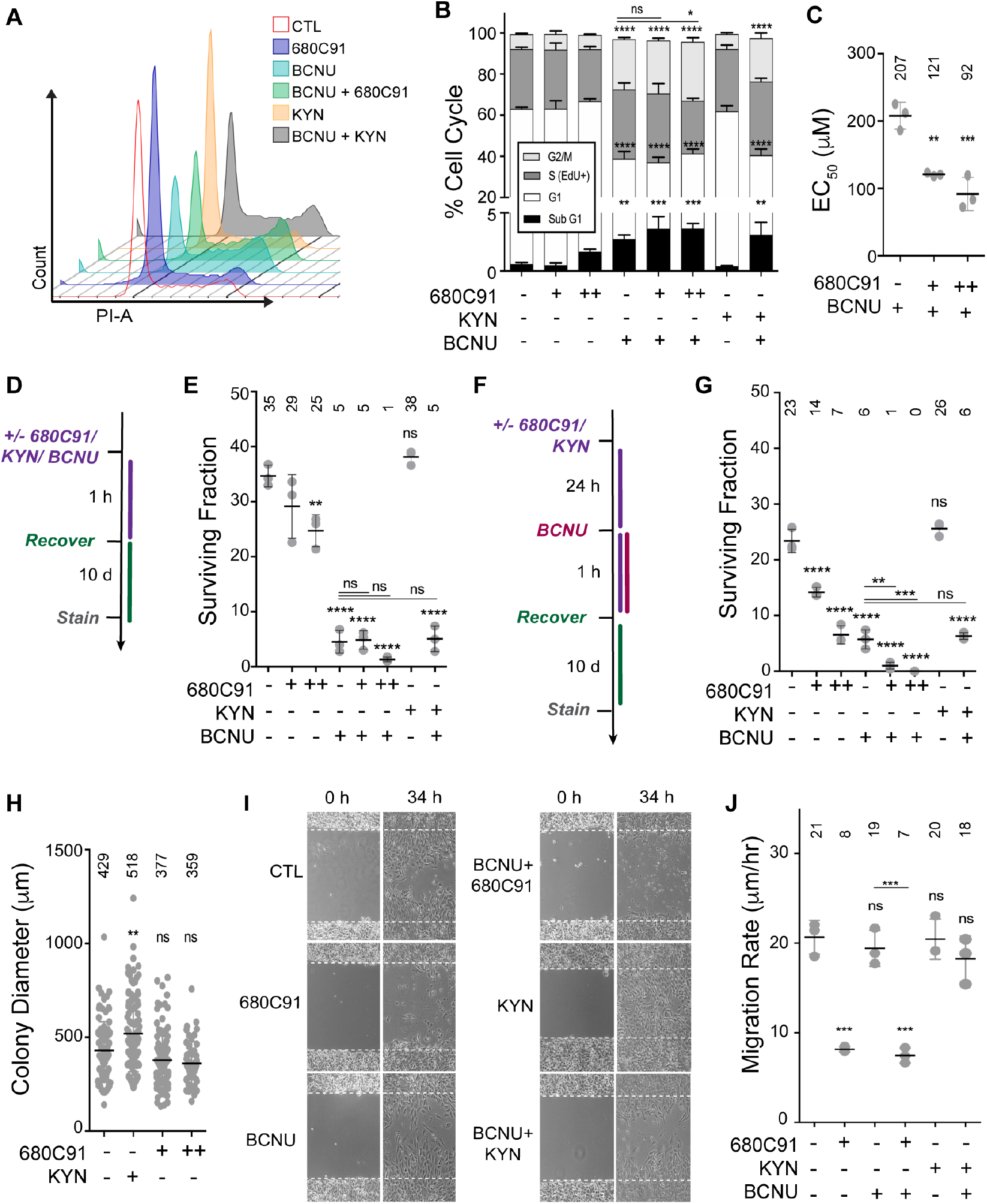
KYN signaling alters cell cycle progression in BCNU-treated cells. *A*, Representative flow cytometry analysis of DNA content for selected experimental conditions depicted in the graph key. *B*, Flow cytometry analysis of cell cycle changes occurring after 680C91/KYN/BCNU treatment and labeling with EdU/PI. *C*, EC_50_ values calculated after dose response experiments were performed with varying concentrations of BCNU with either 10 (+) or 20 μM (++) 680C91. *D*, Treatment scheme for clonogenic assay, where cells were treated 680C91/KYN/BCNU for 1 h and allowed to recover for 10 d. Cells were treated with 680C91 (+, 10 μM; ++, 20 μM), KYN (60 μM), BCNU (125 μM) or a combination of these compounds. *E*, Surviving-fraction calculated by counting colonies composed of at least 25 cells for experimental conditions described in panel D. *F*, Treatment scheme for clonogenic assay where cells were pretreated with 680C91/KYN 24 h prior to 1h BCNU treatment. As with the experiments in panels D and E, cells were treated with 680C91 (+, 10 μM; ++, 20 μM), KYN (60 μM), BCNU (125 μM) or a combination of these compounds. *G*, The surviving fraction was calculated by counting colonies composed of at least 25 cells for each experimental condition described in panel F. *H*, Relative colony diameter measured from all countable colonies for cells treated with 680C91 or KYN alone using conditions described in panel F. *I*, Representative wound migration images showing initial scratch and distance migrated at 34 h after scratch. Cells were treated with 680C91 (20 μM), KYN (60 μM), BCNU (125 μM) or a combination of these compounds. *J*, Wound migration rate was calculated from changes in wound diameter over time for the experimental conditions described in panel I. For all panels, error bars represent the s.d. calculated from three biological replicates. Statistical analysis was calculated using one-way ANOVA with Tukey post-test, where * is P≤ 0.05, ** is P≤ 0.01, *** is P ≤ 0.001 and **** is P ≤ 0.0001.

BCNU treatment produced an increase in the S-phase (EdU^+^) population and a more pronounced increase in the G2/M (4N) populations, indicative of delayed progression through S-phase, as well as G2/M arrest. The EdU^+^ population increased from 29.3% to 33.6% upon exposure to BCNU, whereas the G2/M population increased from 7.2% for the control to 24.5% for BCNU treated cells (**Fig. 3B**). The sub-G1 fraction of cells also increased from 0.6% to 2.7% at this concentration of BCNU (**Fig. 3B**).

Pre-treating cells with 10 μM 680C91 before exposure to BCNU increased the sub-G1 population from 2.7% for BCNU alone to 3.6% for BCNU + 10 μM 680C91 (**Fig. 3B**). This increase corresponded with a small reduction in the 2N (G1) population of cells – from 36.1% for BCNU alone to 33.4% for BCNU + 10 μM 680C91, but this change was not considered statistically significant (*P*-value > 0.05). Pre-treating cells with 20 μM 680C91, on the other hand, led to a statistically significant increase in the G2/M population – from 24.5% for BCNU alone to 28.7% for BCNU + 20 μM 680C91 – and a reduction in the EdU^+^ population – from 33.6% for BCNU alone to 25.6% for BCNU + 20 μM 680C91 – that was also considered statistically significant (**Fig. 3B**). The higher concentration of 680C91 also increased the sub-G1 population from 2.7% for BCNU alone to 3.6% for BNCU + 20 μM 680C91 (**Fig. 3B**) but this change was not considered significant (*P*-value > 0.05). All in all, blocking TDO action corresponded with an enhancement of the anti-proliferative effects induced by BCNU.

Adding exogenous KYN (60 μM, 24 h) did not alter the cell cycle distribution of T98G cells to an extent that reached statistical significance (**Fig. 3A, 3B** and **S1**). There was, however, a slight increase in the S-phase population – compare 33.6% for BCNU alone with 35.8% for BCNU + KYN – and a small decrease in the 4N (G2/M) population – compare 24.5% for BCNU alone with 21.2% for BCNU + KYN (**Fig. 3A, 3B** and **S1**). These modest changes could be related to the fact that exogenous KYN seemed to promote an increase in Chk1 activation and tolerance of RS (see **Fig. 1**), as well as an increased rate of repair (**Fig. 2E**). Still, the overall effects of KP modulation on cell cycle distribution by either 680C91 or KYN were fairly modest, even when BCNU treatment was added.

### Blocking TDO activity reduced the viability and proliferative capacity of glioma-derived cells treated with BCNU

We went on to measure KP-related changes to cell viability following BCNU treatment. We performed dose-response experiments to measure the effect of TDO inhibition on the viability of T98G cells treated with BCNU. Under the conditions used here, we measured an EC_50_ value of approximately 200 μM for T98G cells treated with BCNU alone (**Fig. 3C**). Pre-treating cells with either 10 (+) or 20 μM (++) 680C91 for 24 h reduced the EC_50_ value of BCNU to 125 and 100 μM, respectively (**Fig. 3C**). The results obtained using the Calcein-AM assay were consistent with the notion that blocking KP signaling increased the cytotoxic effects of BCNU-induced DNA damage in GBM-derived cells.

We then examined the effect of KP modulation on the replicative capacity of T98G cells by measuring changes in clonogenic survival (**Fig. 3D**). Treating 500 GBM-derived cells with either 10 μM or 20 μM 680C91 for 1 h reduced clonogenic survival 20% and 30%, respectively (**Fig. 3E**). Treatment with BCNU (125 μM, 1 h) reduced colony formation almost 90% from that of the untreated control (**Fig. 3E**). Cotreating T98G cells with BCNU and 10 μM 680C91 did not alter the proliferative capacity of T98G cells. However, the surviving fraction was reduced another 80% relative to BCNU alone when cells were cotreated with BCNU and 20 μM 680C91 (**Fig. 3E**), indicative of synergy between BCNU-induced DNA damage and KP blockade in the impairment of glioma cell proliferation. Treatment with KYN alone (60 μM, 1 h) increased clonogenic survival slightly, and there was a slight protective effect when cells were cotreated with KYN and BCNU (**Fig. 3E**). However, the KYN-induced changes were not considered statistically significant (*P*-values > 0.05).

Since the effects of TDO inhibition may not be apparent after a one hour exposure to KP modulating agents, we next measured clonogenic survival for T98G cells pre-treated with either 680C91 or KYN for 24 h. We observed a more pronounced decrease in colony formation when T98G cells were exposed to either 10 (+) or 20 μM (++) 680C91 for the additional amount of time (**Fig. 3F**). Treating cells with 10 μM 680C91 reduced the surviving fraction by ~40%, while treatment with 20 μM 680C91 dropped the number of colonies to ~25% of that observed for untreated cells (**Fig. 3G**). Similar to the results with 1 h exposure, adding 125 μM BCNU for 1 h after culturing the cells for an additional 24 h in media reduced clonogenic survival ~75% (compare BCNU alone for **Fig. 3E** and **G**). In contrast to the results obtained with a 1 h cotreatment, a 24 h pre-treatment with 10 μM 680C91 enhanced the anti-proliferative effects of BCNU, reducing clonogenic survival to less than 20% of that observed for BCNU alone, and under the conditions tested here, we did not observe any surviving colonies when T98G cells were pre-treated with 20 μM 680C91 for 24 h and then exposed to 125 μM BCNU for 1 h (**Fig. 3G**), which is again indicative of an enhanced anti-proliferative effect for BCNU when KP signaling is suppressed.

As with the 1 h exposure, the addition of KYN alone (60 μM, 24 h) increased clonogenic survival slightly, and when combined with BCNU treatment, exogenous KYN had a very slight protective effect (**Fig. 3G**). As before, these increases were not statistically significant. Interestingly, we observed a robust increase in the average colony diameter for cells pre-treated with exogenous KYN (**Fig. 3H**). Although not considered statistically significant, there was a corresponding decrease in the average colony diameter for cells pre-treated for 24 h with 680C91 (**Fig. 3H**). In summary, the proliferation of GBM-derived cells over an extended period of time seem to be impacted by modulation of KP signaling and the anti-proliferative effects of BCNU were augmented by blockade of TDO activity.

### TDO inhibition impairs glioma cell motility but does not impact the effects of BCNU on cell migration

Given the previously established role for the KP in promoting malignant properties of gliomas, we were curious to learn whether the combining inhibition of TDO with a genotoxin impacted tumor cell motility. To investigate this possibility, we measured T98G cell migration with the scratch-wound assay (**Fig. 3I**). The rate of cell migration decreased from 20.6 μm/h for control cells to 8.1 μm/h when cells were exposed to 20 μM 680C91 (**Fig. 3J**). Interestingly, BCNU (125 μM) had no impact on T98G cell motility and the effect of combining TDO inhibition with BCNU was minimal, decreasing the migration rate from 8.1 μm/h for 680C91 alone to 7.4 μm/h for the combination treatment (**Fig. 3J**). The addition of exogenous KYN did not have a substantial effect on T98G migration rate either, regardless of whether BCNU was included. The results of the wound migration assay are consistent with the idea that the inhibition of TDO has a more pronounced impact on GBM cell motility than treatment with BCNU (at least under the conditions reported here) and that there is very little difference between 680C91 alone and co-treatment with 680C91 and BCNU.

### Inhibition of TDO resulted in loss of sirtuin signaling and broad changes in genome maintenance

We next employed a quantitative mass spectrometric approach to identify proteomic changes associated with inhibition of TDO and exposure to BCNU (**Fig. 4A**). In an effort to focus on DNA replication/repair factors, we enriched for the nuclear fraction from T98G cells treated with 680C91, BCNU, or a combination of both agents to identify TDO-dependent changes in the nuclear proteome. We performed immunoblotting to confirm successful nuclear enrichment (**Fig. 4A**). We then used the tandem-mass tag (TMT) isobaric labeling approach to quantify changes in abundance between experimental conditions. Each experimental condition was performed in biological quadruplicate. Mass spectrometric analysis was performed and a total of 5787 proteins were identified across all samples (**Table S1**). Changes in protein abundance were considered significant if the FDR adjusted p-value was less than 0.05 in the respective sample groups. We used Qiagen™ Ingenuity Pathway Analysis (IPA) to guide our evaluation of the global changes in cellular pathways (**Tables S2-7**). We also performed manual inspection of the proteomic results to identify changes in individual proteins of interest.

**Figure 4.**
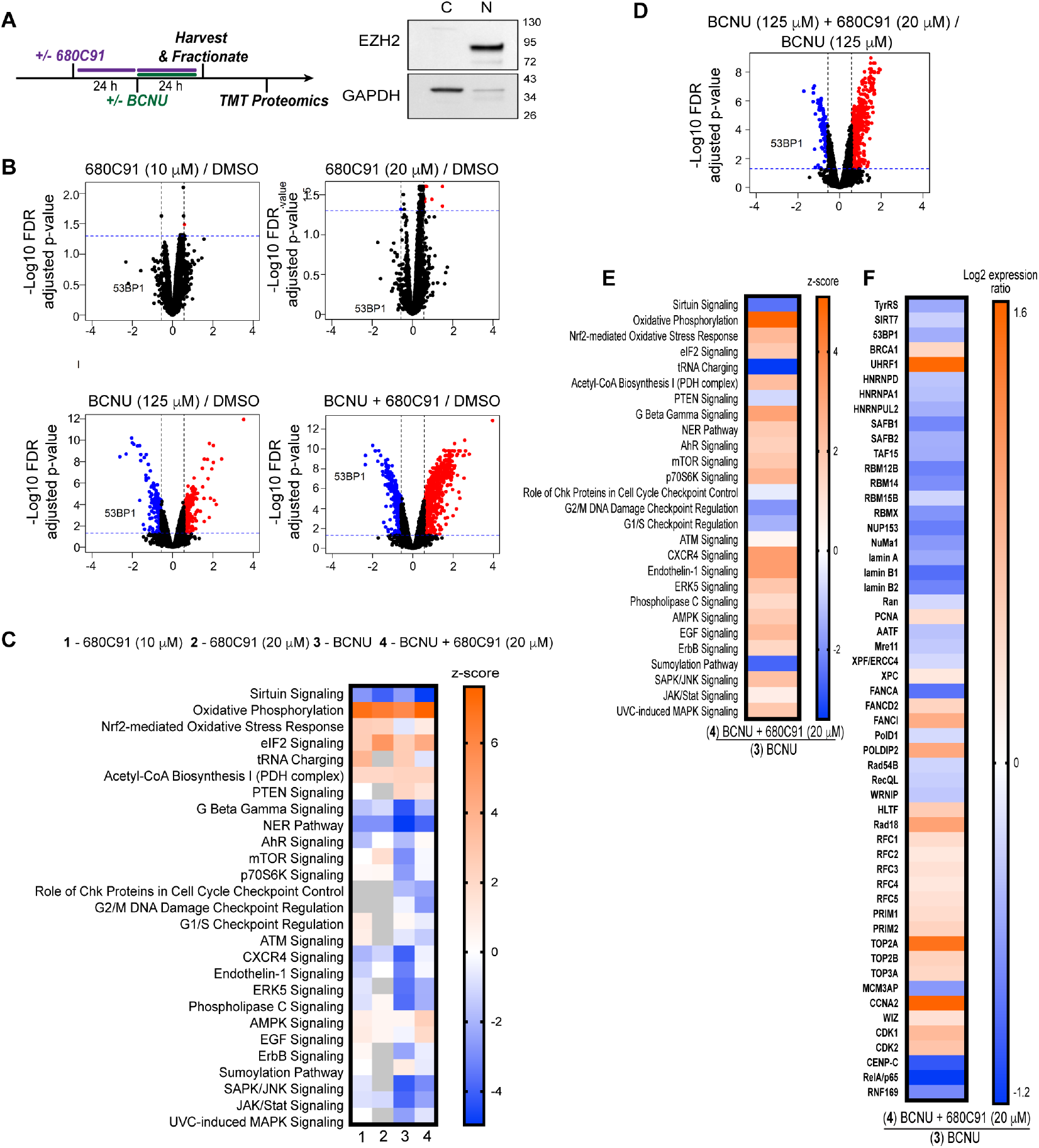
Proteomic analysis of KP blockade in glioma-derived cells shows reduction of centrosome modulators as well as 53BP1. *A*, Cartoon scheme of experimental design for TMT proteomics experiment, and immunoblot depicting nuclear and cytoplasmic fractionation; where EZH2 was used as a nuclear marker and GAPDH was used as a cytoplasmic marker. *B*, Volcano plots representing the limma log_2_ fold change and FDR adjusted p-value results from the following conditions; CTL vs 680C91 (10 and 20 μM), CTL vs BCNU, and CTL vs BCNU+ 680C91. Proteins up-regulated in the first group of the comparison with at least a fold change > 1.5 are highlighted in red while down-regulated proteins are in blue. Protein 53BP1 is highlighted in each of the different comparisons. The horizontal blue dotted line indicates a p-value threshold of 0.05 and the vertical black dotted lines indicate a fold change of 1.5 and −1.5. *C*, Heat map of cellular pathway changes (Z-scores) predicted from comparison of the proteomic results for the following conditions to the DMSO control sample: 1. 680C91 (10 μM), 2. 680C91 (20 μM), 3. BCNU, 4. BCNU+ 680C91 (20 μM) relative to the control. A Z-score of 2 or greater shows upregulation (orange), while −2 or less was considered downregulation of pathway activity (blue). *D*, A volcano plot showing changes in protein abundance between the BCNU vs BCNU+ 680C91 (20 μM) conditions. The decrease in 53BP1 abundance is highlighted. *E*, Heat map of cellular pathway comparison of the BCNU alone sample with the proteomic results obtained for the BCNU+ 680C91 (20 μM) sample. *F*, Heat map showing Log_2_ fold-change in the abundance of individual proteins when comparing the BCNU alone sample with the BCNU+ 680C91 (20 μM) sample.

We first compared the difference between DMSO treated cells and cells exposed to 680C91 (10 and 20 μM, 24 h). Overall changes in abundance were modest for individual proteins in both treatment conditions (**Fig. 4B**), but there were some interesting trends identified at the pathway level (**Fig. 4C**). AhR signaling was predicted to be diminished by treatment with 10 μM 680C91 (**Table S2**) but the change at the pathway level was ambiguous for cells treated with 20 μM 680C91 (**Table S3**). Both concentrations of 680C91 led to a reduction in sirtuin signaling (10 μM: z-score = −2.71, p-value = 1.58 x 10^-17^; 20 μM: z-score = −4.00, p-value = 1 x 10^-12^). Treatment with 10 μM 680C91 reduced the abundance of nuclear SIRT6- and SIRT7-related targets GPAA1, MRPL16, MRPS31, MRPS33, and RPS3 (**Table S2**). At 20 μM 680C91, a change in SIRT7 was again identified at the pathway level (**Table S3**), although the directionality of the effect was ambiguous.

Sirtuins depend on available stores of NAD^+^ to effectively catalyze deacetylation of a wide range of protein targets (18), and glioma cells use KYN-derived quinolinic acid (QA) to replenish NAD^+^ stores, which protects tumor cells from genotoxic agents (19). NAD^+^ levels also support maintenance of genomic integrity. The protection against DNA damage afforded by sustained NAD^+^ levels is related in part to adequate substrate availability for the DNA repair mediator poly[ADP-ribose]polymerase 1 (PARP-1) with simultaneous disruption of NAD^+^ biosynthesis and base excision repair (BER) sensitizing glioma cells to temozolomide (TMZ) (20). To support the IPA of the proteomics results, we probed for total lysine acetylation via immunoblotting to assess global changes in sirtuin action in response to TDO inhibition (**Fig. S3A**). We also checked for total PARylation levels (**Fig. S3B**). Consistent with NAD^+^ depletion and diminished deacetylase activity, we observed a pronounced increase in total acetylated lysine, including acetylated histones, when GBM cells were treated with 680C91 (**Fig. S3A**). Treatment with 680C91 also produced a concomitant decrease in global PARylation (**Fig. S3B**), indirectly suggestive of diminished NAD^+^ stores in cells.

We were intrigued by the identification of SIRT7 specifically because of its multifaceted role in regulating chromatin condensation, DNA repair dynamics, tolerance of endoplasmic reticulum (ER) stress, and mitochondrial homeostasis (21–26). One function of SIRT7 is to promote tolerance of ER stress by suppressing Myc activity and silencing expression of ribosomal proteins (22). Consistent with loss of SIRT7 function, we identified Myc as one of the top upstream regulatory factors activated by inhibition of TDO (10 μM: z-score = 5.68, p-value = 2.64 x 10^-18^; 20 μM: z-score = 7.11, p-value = 1.61 x 10^-15^) and the abundance of ribosomal proteins was increased in cells treated with both concentrations of 680C91 (**Tables S2-3**). Accordingly, IPA identified increased activation of eukaryotic initiation factor 2 (eIF2) translational control in cells treated with the TDO inhibitor (10 μM: z-score = 5.24, p-value = 1 x 10^-67^; 20 μM: z-score = 2.56, p-value = 1 x 10^-17^). At the higher concentration of 680C91, an alteration in mammalian Target of Rapamycin (mTOR) was noted at the pathway level (z-score = 0.83, p-value = 2.5 x 10^-36^;), and inhibition of RICTOR, a component of mTOR complex 2 (mTORC2), was identified as either the top or one of the top alterations to upstream regulators at both concentrations of the TDO inhibitor (10 μM: z-score = −7.08, p-value = 8.4 x 10^-14^; 20 μM: z-score = −7.94, p-value = 6.5 x 10^-31^). Accordingly, we observed a decrease in phosphorylation of the mTOR2 target pS473 Akt when T98G cells were treated with 680C91 (**Fig. S3C**).

Consistent with the energetic demands of protein synthesis, increased oxidative phosphorylation (OXPHOS) was identified at the pathway level for both concentrations of 680C91 (**Fig. 4B** and **C**; 10 μM: z-score = 7.00, p-value = 1.0 x 10^-29^; 20 μM: z-score = 6.48, p-value = 2.0 x 10^-29^). An increase in pyruvate dehydrogenase (PDH)-mediated acetyl-CoA biosynthesis was also observed signifying increased flux through the TCA cycle, which could be related to diminished production of the allosteric PDH inhibitor acetyl-CoA from KYN breakdown (27). Inhibition of TDO also increased the Nrf2-mediated oxidative stress response at both concentrations of 680C91 (10 μM: z-score = 2.50, p-value = 0.003; 20 μM: z-score = 2.33, p-value = 0.007). The Nrf2 oxidative stress response helps to relieve ER stress and protect cancer cells to reactive oxygen species (ROS) through the promotion of anti-oxidant gene expression (28).

Inhibition of TDO resulted in increased X-box binding protein-1 (XBP1), a transcription factor and key mediator of the unfolded protein response (UPR) that functions downstream of eIF2 (29). XBP1 function is fine-tuned by the balance between p300 acetylation and sirtuin-catalyzed deacetylation with loss of sirtuin activity leading to higher expression of the active spliced form of XBP1 (30). Nuclear XBP1 levels were increased in cells treated with 10 μM 680C91 (log_2_FC = 0.96, p-value = 0.009) and XBP1 regulated targets were increased in cells treated with both concentrations of 680C91 (10 μM: z-score = 7.33, p-value = 1.59 x 10^-30^; 20 μM: z-score = 4.9, p-value = 1.2 x 10^-10^). These results are consistent with the notion that inhibition of the KP in T98G cells increased the relative level of UPR activation from protein synthesis.

Analyzing the proteomics results further, we noticed that EGFR signaling was predicted to be activated in cells treated with 680C91 (10 μM: z-score = 2.29, p-value = 0.00039; 20 μM: z-score = 2.85, p-value = 0.0014). At the higher concentration of inhibitor, there was a decrease in the abundance of the EGFR-associated protein RanBP6 and its downstream effector STAT3 (**Table S3**). RanBP6 regulates nuclear import of EGFR and STAT3, and RanBP6 silencing has been shown to increase glioma growth *in vivo* (31). STAT3 is a transcription factor that is phosphorylated by Janus kinases (JAK) to transduce cytokine signaling to the nucleus (32). Based on our proteomics results, blocking TDO activity might activate EGFR signaling by downregulating RanBP6 and by inhibiting STAT3-mediated signaling.

In addition to alterations in the metabolic status, TDO inhibition resulted in a depletion of nuclear proteins directly involved in NER (nucleotide excision repair) and double-strand break repair (DSBR), including ATM, Mre11, Nbs1, SMARCAL1/2/4, WRN, and a number of chromatin remodeling enzymes. At the pathway level, loss of NER-related factors was scored as more significant, but inhibition of DSBR was also considered to be of importance based on IPA (**Tables S2-S3**). Overall, TDO inhibition in T98G cells seemed to promote a reduction in the nuclear abundance of factors associated with the DNA damage response.

We next analyzed proteome-level changes in T98G cells exposed to the DNA alkylating/crosslinking agent BCNU (**Table S4**). The concentration of BCNU we used for the proteomic analyses was 125 μM, which is below the EC_50_ value of ~200 μM that we measured for T98G cells but high enough to induce a response to DNA damage. Treating GBM-derived cells with BCNU led to activation of phosphatase and tensin homolog on chromosome 10 (PTEN), a negative regulator of PI3K/Akt/mTOR signaling (z-score = 2.11, p-value = 0.000093). These findings were further corroborated by decreased pS473 Akt in cells treated with BCNU (**Fig. S3C**). In addition to regulation of the PI3K/Akt/mTOR cascade, PTEN physically associates with centromeres to protect them from breakage and loss of PTEN leads to defects in HR through failed recruitment of Rad51 to sites of damage (33, 34). BCNU treatment resulted in a decrease in mTOR and eIF4/p70 S6K signaling (**Fig. 4C**), consistent with PTEN activation and indicative of an overall reduction in protein synthesis.

BCNU treatment also resulted in a predicted down-regulation of SIRT6 and SIRT7 signaling (**Table S4**), even below that observed for cells treated with 680C91 alone (**Table S7**). There was an overall increase in acetylated lysine by immunoblotting (**Fig. S3A**). However, the level of histone acetylation appeared to diminish in response to BCNU (**Fig. S3A**, see band near 17 kDa marker). BCNU-induced hypoacetylation of histone H3 has been reported previously for glioma-derived cells (35). The effect of BCNU on sirtuin signaling may be related to increased consumption of NAD^+^ by PARP-1. Indeed, we observed elevated PARylation levels accompanied BCNU treatment (**Fig. S3B**). NER and BRCA1-related DNA repair factors were also diminished by treatment with BCNU (**Table S4**), perhaps owing to the attenuation of protein synthesis. The abundance of nuclear localized DNA repair factors was lower in BCNU-treated cells than in cells treated with 20 μM 680C91 (**Table S7**, NER: z-score = 3.43, p-value = 2.5 x 10^-23^; BRCA1 DDR: z-score = 2.67, p-value = 1.58 x 10^-10^). Another difference between BCNU-treated cells and cells treated with 680C91 was that treatment with BCNU led to a slightly diminished Nrf2 antioxidant response (z-score = – 0.66, p-value = 2 x 10^-15^) and a slight increase in SUMOylation (z-score = 0.96, p-value = 2.0 x 10^-13^).

Several interesting changes in the nuclear-enriched proteome were identified when we compared the results for cells treated with BCNU alone to those obtained with cells treated with the TDO inhibitor 680C91 (20 μM) prior to BCNU exposure (**Fig. 4D** and **E**, **Table S7**). At the pathway level, the most significant change identified was a decrease in tRNA charging (z-score = −3.4, p-value = 5 x 10^-18^). Based on quantitative proteomics, multiple tRNA synthetases, including Tyrosyl-tRNA synthetase (TyrRS), were depleted by co-treatment with 680C91 and BCNU relative to treatment with BCNU alone. This observation was interesting given that nuclear localized TyrRS was reported to upregulate expression of DNA repair factors, such as BRCA1 and RAD51, in response to oxidative stress through a direct interaction with the E2F1 transcription factor (36). Cells that cannot import TyrRS exhibit higher levels of γH2AX following treatment with H_2_O_2_ (36, 37).

Compared to treatment with BCNU alone, OXPHOS, Nrf2, acetyl-CoA biosynthesis, and eIF2 pathways were elevated by the pre-treatment with 680C91 prior to BCNU exposure (**Fig. 4E**), indicative of sustained energetic demands similar to what we observed for treatment with 680C91 alone. Combining TDO inhibition with BCNU resulted in diminished sirtuin signaling relative to BCNU alone (z-score = – 2.46, p-value = 5 x 10^-11^). While total lysine acetylation for the combined treatment did not change much compared to BCNU alone, there was a slight increase in histone acetylation (**Fig. S3A**), which could signal diminished deacetylase action on chromatin in GBM cells with suppressed KP are exposed to BCNU.

Once again, SIRT7 was singled out as the major sirtuin family member regulating multiple proteins identified in our analysis (z-score = −2.02, p-value = 1.8 x 10^-5^). Nuclear abundance of SIRT7 was decreased by the combination treatment, as compared to BCNU treatment alone (**Fig. 4F**, log_2_FC = −0.32, p-value = 0.0024). As before, we observed activation of the sirtuin-regulated UPR mediator XBP1 (z-score = 5.48, p-value = 5.9 x 10^-9^). EGFR signaling was also elevated, as we observed with treatment with 680C91 alone (z-score = 2.33, p-value = 0.037), with EGFR abundance increasing slightly (log_2_FC = 0.44, p-value = 0.002). RICTOR-associated mTORC2 was predicted to be down-regulated in cells exposed to 680C91 and BCNU compared to BCNU alone (z-score = −2.64, p-value = 1.8 x 10^-5^). This prediction was further supported by lower pS473 Akt in T98G cells exposed to the combination treatment compared to BCNU alone (**Fig. S3C**), as well as a predicted activation of FOXO1 (z-score = 3.16, p-value = 8.0 x 10^-9^), which is normally suppressed by mTORC2.

Further interrogation of the proteomics data allowed us to identify additional changes in DSBR that extend beyond SIRT7-mediated effects. Some of these changes were consistent with the idea that loss of TDO activity results in a diminished capacity to repair BCNU-induced DNA damage. For example, there was a decrease in nuclear abundance of 53BP1 in cells treated with the combination of 680C91 and BCNU compared to treatment with BCNU alone (**Fig. 4F**). 53BP1 is a critical factor in repair of DSBs (38). Consistent with diminished 53BP1, there was an increase in BRCA1 and the associated E3 ubiquitin ligase UHRF1, which together promote K63-linked poly-ubiquitinylation of the 53BP1-binding partner Rif1 and suppress NHEJ (39). Furthermore, there was a broad reduction in proteins known to regulate repair of DSBs in a poly[ADP-ribose] (PAR)-dependent manner through interactions involving low complexity domains (LCDs). The PAR-dependent accumulation of LCD proteins induces liquid-liquid phase separation (i.e., biomolecular condensates) near sites of DNA damage (40). There was an uncanny similarity between the list of LCD-containing proteins depleted in 680C91/BNCU-treated cells and those previously implicated in PAR-mediated regulation of DNA repair through liquid-liquid demixing (40–43). This list included HNRNPD, HNRNPA1, HNRNPUL2, SAFB1, SAFB2, TAF15, RBM12B, RBM14, RBM15B, RBMX, and others (**Fig. 4F**). PAR-initiated liquid-liquid demixing nucleates self-assembling structures that may regulate DNA repair choice by filtering which factors gain access to sites of damage. In our experiments, the decreased nuclear abundance of these LCD-containing proteins may be related to limited PARP activity resulting from inadequate NAD^+^ stores, which could prevent effective liquid demixing and proper coordination of DNA repair.

The down-regulation of factors that control 53BP1 trafficking was also noted. Nucleoporin 153 (NUP153) was depleted, as was the nuclear structural protein NuMa 1 (**Fig. 4F**). NuMa1 was recently shown to control diffusion of 53BP1 (44). More specifically, NuMa1 was shown to reduce 53BP1 motility outside of DNA repair foci with no effect on total 53BP1 levels. Multiple lines of evidence support the notion that NUP153 is a key regulator of 53BP1 nuclear entry (45–48). The mechanism of NUP153-mediated import relies on the intermediate filament protein lamin A. Mislocalization of NUP153 in response to diminished lamin A (i.e., elevated levels of the lamin A precursor) leads to decreased localization of Ran, a GTPase responsible for nuclear import, and impeded nuclear entry of large protein cargo, such as 53BP1, but not smaller cargo like PCNA (47). The abundance of lamin A and Ran levels were found to be less in the cells treated with a combination of 680C91 and BCNU than in cells treated with BCNU alone (**Fig. 4F**). In line with this model, nuclear PCNA levels increased slightly when TDO inhibition was combined with BCNU (**Fig. 4F**). Analyzing a plot of the fold-change in protein abundance as a function of molecular weight produced a Pearson r value of −0.18 (p-value < 0.0001, **Fig. S2**), indicative of a modest negative effect on nuclear abundance for higher molecular weight proteins in cells treated with 680C91 and BCNU. It may be that down-regulation of KP signaling impairs nuclear entry for large cargo in response to BCNU treatment. This notion fits with a decrease in the nuclear abundance of 53BP1, a protein of >200 kDa, as well as decreased abundance for several other key repair factors (e.g., AATF, Mre11, ERCC4, FANCA, PolD1, Rad54B, RecQL, WRNIP). With that said, there were also increases in some high molecular weight repair factors, including FANCD2, FANCI, HLTF, and Rad18, which paints a more complicated picture of how DNA repair proteins are transported in GBM cells with an attenuated KP signaling cascade.

Another interesting difference between the BCNU and BCNU/680C91 treated samples was an overall increase in DNA replication-associated proteins when TDO activity was inhibited. In addition to PCNA, we observed increase in the relative abundance of RFC1-5, POLD1, PRIM1, PRIM2, and TOP2A (**Fig. 4F**). Slight increases in MCM2-7 were also apparent. There was a notable depletion of MCM3AP, which acetylates MCM3 and inhibits replication initiation (49). Mutations in MCM3AP have been shown to result in defective HR-mediated repair of DSBs (50), and failed activation of canonical NF-κB signaling caused by MCM3AP mutation may be responsible for the defect in HR repair. Such a scenario is consistent with the break repair defects we observed for TDO-deficient cells exposed to BCNU (**Fig. 2**).

There was >three-fold increase in cyclin A2 (CCNA2) in cells treated with 680C91 prior to BCNU exposure (**Fig. 4F**). A recent proteomics study identified cyclin A2 as one of the top PCNA-interactors following treatment with camptothecin (CPT) (51). The same study identified widely interspaced zinc finger (WIZ) as a PCNA-interacting partner. The authors speculated that WIZ might aid in the recruitment of the G9a-GLP methyltransferase. The action of the G9a-GLP methyltransferase facilitates recruitment of the BRCA1-associated E3 ligase UHRF1. UHRF1 is also an essential factor in the maintenance of DNA methylation (52). In accordance with these previous studies, we observed a small increase in nuclear WIZ and an approximate 3-fold increase in nuclear UHRF1 when we compared cells treated with BCNU alone to those treated with both 680C91 and BCNU (**Fig. 4F**).

We observed an increase in both CDK1 and CDK2 in the co-treated cells, again consistent with a relative acceleration in the replication program of TDO-deficient cells damaged with BCNU. Cyclin A2-CDK1 promotes origin firing, S-phase progression and mitotic entry (53). Cyclin A2 also helps coordinate mitotic entry through an interaction with CDK2 that promotes activation of the Anaphase Promoting Complex/Cyclosome (APC/C) (54). The overall increase in replication factors was coupled with important defects in DNA repair capacity noted above (e.g., loss of 53BP1, dysregulation of factors involved in PAR-initiated liquid-liquid demixing). These results may provide clues to understanding the elevated DNA damage and CIN observed in the cells co-treated with 680C91 and BCNU.

In addition to changes in cell cycle regulators, there was a decrease in PTEN signaling in 680C91/BCNU treated cells relative to BCNU treatment alone (**Fig. 4E**). PTEN plays an essential role in maintaining chromosomal integrity through interactions with CENP-C, a component of the kinetochore (33, 34). There was an approximate 2-fold decrease in CENP-C levels in 680C91/BCNU cells relative to treatment with BCNU alone (**Fig. 4F**). PTEN deficiency has been shown to not only inhibit DNA repair but also to impair cell cycle checkpoint activation and promote genomic instability, especially chromosomal damage. CIN can take many forms (55, 56). It is possible that the increased CIN observed in the MN assay for cells co-treated with 680C91 and BCNU was related in part to defects in centromere protection induced by down-regulation of PTEN in conjunction with DNA damage.

Analysis of the proteomics results revealed some interesting alterations in proteins involved in NF-κB-mediated responses to DNA damage. ATM is known to communicate genomic stress through a signaling cascade that involves the NF-κB transcription factor (57–59). NF-κB stimulates HR by accelerating RPA and Rad51 foci formation (57). Blocking TDO activity resulted in a depletion of the RelA/p65 NF-κB subunit, as well as the NF-κB activating kinase IKK-beta (IKBKB) in 680C91/BCNU-treated cells (**Fig. 4F**). We also observed loss of the NF-κB signal transducer Bcl2-associated transcription factor 1 (Bclaf1), which is upregulated through the ATM/Nemo/NF-κB axis in response to doxorubicin-induced senescence (60).

There was a decrease in nuclear TRAF6 and IFI16 abundance in TDO-deficient cells treated with BCNU, which we found interesting in light of a recent study that implicated the DNA binding protein IFI16 and the TRAF6 E3 ubiquitin ligase in activation of the DNA sensing adaptor STING (58). As with the LCD-mediated liquid-liquid mixing, this cascade is dependent upon PARP1 activity. STING activation is a key event in the induction of NF-κB transcriptional response to DNA damage and depletion of TRAF6 and IFI16 in TDO-deficient cells treated with BCNU would presumably lead to a reduction in STING/NF-κB signaling. Taken together, the loss of nuclear p65, as well as depletion of other NF-κB-related factors, provided evidence for dysregulation of ATM signaling and ultimately failed repair of BCNU-induced DNA damage in glioma-derived cells that lack TDO activity.

### Inhibition of TDO impairs 53BP1 recruitment to sites of DNA damage

The results of the proteomics experiments allowed us to identify a number of changes in strand-break repair pathways that seemed to be regulated to some degree by TDO activity. A key finding from our proteomic analysis was that nuclear 53BP1 levels were diminished in cells co-treated with 680C91 and BCNU (**Fig. 4D** and **F**). The 53BP1 protein serves as an important regulator of the partitioning between HR and NHEJ double-strand break repair pathways – with high-levels of 53BP1 favoring NHEJ over HR. We postulated that the sustained levels of BCNU-induced breaks observed with the comet assay for cells treated with the TDO inhibitor might be related to defective recruitment of 53BP1 to sites of DNA damage. To test this idea and help validate our proteomic results, we measured changes in 53BP1 foci formation via IF microscopy. Pre-extraction of cytosolic proteins and co-staining for DAPI ensured that the signal was from chromatin-bound 53BP1 (**Fig. 5A**). Chromatin bound gH2AX was used as a proxy for how much DNA damage signal was present. We quantified foci formation for 53BP1 but the pan-nuclear signal observed for some conditions prevented an accurate assessment of gH2AX foci. For that reason, we report gH2AX signal intensity per cell. The experimental design was identical to that used for the comet assay, with cells being exposed to treatment with either DMSO, 680C91 (20 μM), or kynurenine (60 μM) for 24 h prior to co-treatment with BCNU for another 24 h.

**Figure 5.**
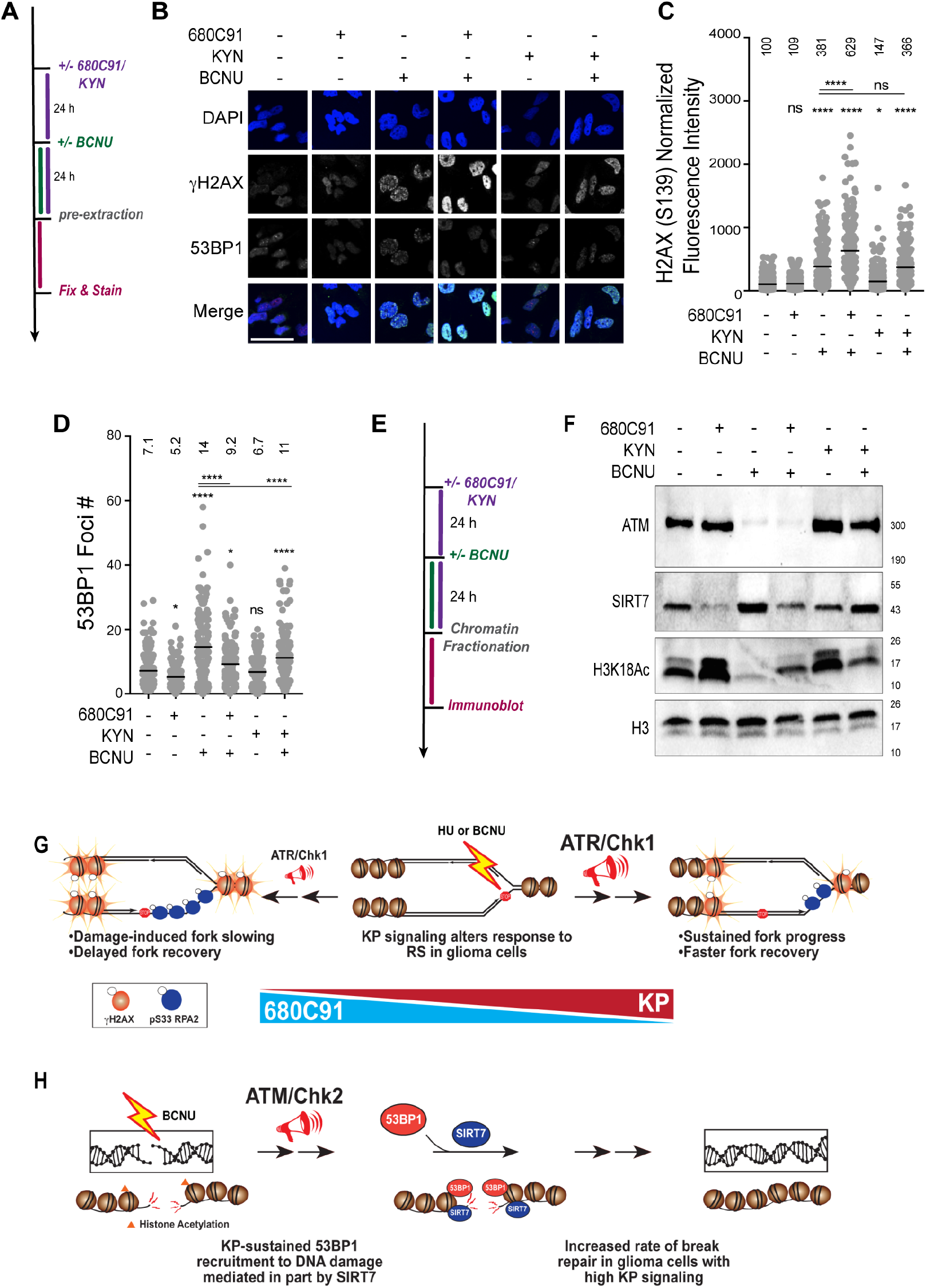
TDO inhibition reduces ability to respond to damage via decreased recruitment of 53BP1 to sites of DNA damage. *A*, Cartoon schematic of experimental design for experiments monitoring changes in chromatin-bound 53BP1 and γH2AX. *B*, Representative immunofluorescent images of γH2AX and 53BP1 stained cells after the treatment conditions listed in (A). Scale bar: 50 μm. *C*, Plot of γH2AX normalized signal intensity. The mean intensity value is shown above the data points. *D*, Graph of 53BP1 foci number per cell, where mean foci number is listed above each treatment. For both (C and D) at least 140 cells were quantified from two biological replicates. A one-way ANOVA was used for statistical analysis where * is P≤ 0.05, ** is P≤ 0.01, *** is P ≤ 0.001 and **** is P ≤ 0.0001. E, Cartoon schematic of experimental design for chromatin fractionation experiments. *F*, Immunoblots are shown for chromatin bound ATM, ATR, SIRT7 and H3K18Ac. Histone H3 was used as a loading control. Cells were treated with 680C91 (20 μM), KYN (60 μM), BCNU (125 μM) or a combination of these compounds as described in the Materials and Methods section. *G*, A model illustrating the proposed effects of KP signaling on the RSR. Suppression of KP signaling via 680C91 treatment led to reduced replication rate and delayed replication stress associated fork recovery. High KP activity resulted in better RS tolerance and elevated fork recovery in response to genotoxic agents. H, A model depicting the relative impact of KP signaling on repair of BCNU-induced DNA breaks in glioma cells. Activation of KP aided in the recruitment of 53BP1 to sites of damage, in part, through SIRT7-mediated deacetylation of histone H3K18. Inhibition of TDO impaired SIRT7 and 53BP1 recruitment to chromatin and resulted in delayed break repair, while stimulation of the KP with exogenous KYN increased the rate of break repair. Shuttling ATM on and off of chromatin may also play a role in the TDO-dependent effects observed for repair of BCNU-induced strand breakage.

In contrast to the reduction in gH2AX measured by immunoblotting and IF microscopy of total nuclear protein (**Fig. 1** and **2**), there was effectively no change in chromatin-bound gH2AX signal intensity for cells exposed to 680C91 (**Fig. 5B** and **C**). Treatment with 680C91 alone reduced the number of 53BP1 foci ~15% relative to control (**Fig. 5B** and **D**). As expected, the addition of BCNU increased the number of the chromatin-bound gH2AX signal more than 3-fold (**Fig. 5C**) and 53BP1 foci ~2-fold (**Fig. 5D**), indicative of DDR activation. Pre-treating cells with 680C91 led to diminished 53BP1 foci formation in response to BCNU exposure (**Fig. 5D**) in spite of the fact that gH2AX signal intensity soared ~40% above the value measured for cells treated with BCNU alone (**Fig. 5C**). These results validated an important conclusion derived from our proteomics experiments – loss of TDO activity impairs the ability of GBM cells to successfully recruit 53BP1 to sites of DNA damage.

Treating cells with exogenous KYN produced a small increase the gH2AX signal relative to control (**Fig. 5C**) but did not change the basal level of 53BP1 foci (**Fig. 5D**). Combining KYN with BCNU reduced 53BP1 foci relative to BCNU alone (**Fig. 5D**), but there was also a slight reduction in gH2AX signal intensity for KYN-treated cells exposed to BCNU compared to treatment with BCNU alone (**Fig. 5C**). In other words, adding exogenous KYN reduced the amount of BCNU-induced gH2AX signal (likely through more efficient repair based on the comet assay results), which may have expedited removal of chromatin-bound 53BP1. By way of comparison, combining BCNU treatment with TDO inhibition led to higher gH2AX signal, fewer 53BP1 foci, and increased strand break formation than cells treated with BCNU alone.

### Chromatin-bound ATM and SIRT7 are altered by changes in KP signaling

Others have shown that SIRT7 activity promotes 53BP1 recruitment through deacetylation of histone H3K18 (61). One study reported that SIRT7-catalyzed deacetylation was required to dissociate ATM from chromatin and completion of DSBR (23). Failure to deacetylate ATM led to increased levels of chromatin-bound 53BP1 and RPA2 (23). Given that SIRT7 was identified in our proteomics analysis as being altered by modulation of the KP, we decided to investigate whether the impaired recruitment of 53BP1 could be related to changes in SIRT7 action (**Fig. 5E**).

First, we probed for KP-dependent changes in chromatin-bound SIRT7 and ATM. A sharp decrease in chromatin-bound SIRT7 was observed for GBM cells treated with 680C91 (**Fig. 5F**). The decrease in SIRT7 produced by treatment with 680C91 was accompanied by an increase in H3K18Ac but very little change in ATM bound to chromatin (**Fig. 5F**). Again, the inverse correlation between SIRT7 localization and H3K18Ac was expected due to loss of the deacetylase action of SIRT7 on chromatin. These results were consistent with our proteomics analysis and the idea that TDO inhibition exerts downstream effects on SIRT7 localization and activity.

Comparing BCNU exposure with or without TDO inhibition further supported our working hypothesis, as blockade of TDO activity impaired SIRT7 recruitment to DNA following exposure to BCNU relative to treatment with the genotoxin alone (**Fig. 5F**). H3K18Ac dropped sharply with BCNU treatment, consistent with more chromatin-bound SIRT7 (**Fig. 5F**). Pre-treatment with 680C91 increased the amount of H3K18Ac present following exposure to BCNU – consistent with a reduction in chromatin-bound SIRT7 compared to cells treated with BCNU alone. The amount of chromatin-bound ATM was sharply reduced for cells treated with BCNU whether TDO was inhibited or not (**Fig. 5F**).

The addition of exogenous KYN increased the amount of ATM bound to DNA without much of an effect on SIRT7 levels relative to DMSO-treated cells (**Fig. 5F**). Addition of KYN did not seem to increase SIRT7 activity, however, as the amount of H3K18Ac was at least as much as DMSO-treated cells. Adding exogenous KYN led to a dramatic increase in chromatin-bound ATM following exposure to BCNU (**Fig. 5F**, compare ATM signal for lane 3, BCNU alone to lane 6, + KYN + BCNU). SIRT7 remained bound to chromatin when KYN-fed cells were exposed to BCNU (**Fig. 5F**) - contrasting sharply with the depletion of SIRT7 that accompanied TDO inhibition. The activity of SIRT7 in cells treated with KYN and BCNU seemed to be slightly less robust than that observed for treatment with BCNU alone, based on H3K18Ac levels (**Fig. 5F**), but H3K18Ac was reduced in cells treated with KYN and BCNU relative to treatment with KYN alone (**Fig. 5F**, compare last two lanes for H3K18Ac blot). In summary, these experiments supported the notion that inhibition of TDO produced changes in SIRT7 activity on DNA that likely influenced the dynamics of break repair, including ATM and 53BP1 recruitment to sites of damage. The increase in chromatin-bound ATM afforded by exogenous KYN matches nicely with the faster kinetics of break repair observed with the comet assay. The relative increase in chromatin-bound SIRT7 for cells with an adequate KYN supply may allow for more effective shuttling of ATM and completion of DSBR than what occurs in GBM cells lacking TDO activity.

## DISCUSSION

Understanding factors that influence the robust RSR and DNA repair capacity of gliomas is an important part of developing new and more effective treatment strategies. We have investigated the relationship between TDO activity and the maintenance of genomic integrity in glioma-derived cells - uncovering evidence that KP signaling has a broad effect on the ability of glioma-derived cells to respond to HU-induced RS and damage generated by BCNU, a DNA alkylating agent used in the treatment of malignant brain tumors. The impact of KP signaling on central nervous system disorders, immune function, and tumor biology is known to be multi-faceted, involving interplay between transcriptional regulators, NAD^+^-dependent activities, mitochondrial function, heme biosynthesis, and energy utilization circuits (27, 62–65). Our findings provide some new and intriguing insights into how metabolic changes in the tumor microenvironment can influence genomic stability in glioma cells and possibly influence responsiveness to chemotherapy.

KP activation by tumor-specific up-regulation of TDO is an important determinant of glioma malignancy and the suppression of anti-tumor immune responses (9). These effects appear to be mediated in part by the AhR, a transcription factor that exerts ligand–dependent and ligand-independent effects (66). The AhR regulates the expression of a multitude of factors, including the TLS enzyme hpol κ (13, 67). In addition to transcriptional regulation, nuclear AhR has been shown to physically interact with DNA repair factors, including gH2AX, DNA-PK, ATM, and Lamin A (68), and reducing AhR levels inhibits repair of ionizing radiation (IR)-induced DNA damage (68). KP signaling may also fuel DNA repair through *de novo* synthesis of the PARP substrate NAD^+^ (20, 62, 69). We became interested in the role of KP signaling because of its potential relationship to hpol κ, an enzyme implicated in chemoresistance and poor outcomes in glioma patients. The regulation of hpol κ in response to AhR activation protects cells from bioactivated carcinogens like benzo[*α*]pyrene (70, 71). However, up-regulation of hpol κ has a negative impact on glioma patient prognosis and response to treatment (17, 72), and under the right circumstances, high levels of hpol κ can have a negative impact on fork progression and genome stability (73, 74). Previously, we reported that blockade of the KP-AhR axis reduced hpol κ expression and MN formation in GBM-derived cells (13). Our earlier study led us to hypothesize that blocking KP signaling would alter the responsiveness of GBM-derived cells to RS, leaving them more susceptible to acute DNA damage from an exogenous agent.

Our initial focus on DNA replication dynamics allowed us to explore the idea that elevated KP signaling could influence fork progress and DNA damage tolerance by modulating the RSR. Analysis of replication rates using the DFS assay revealed that loss of TDO activity led to impaired fork progress following treatment with BCNU and less effective fork restart in response to HU treatment. The defects in fork restart could be related to the fact that TDO inhibition down-regulated RSR factors, such as hpol κ and phosphorylated Chk1. While there are some inconsistencies in the literature, there is reasonable evidence to suggest that hpol κ can directly influence Rad17 and 9-1-1 complex-mediated recruitment to sites of replication stress (75). In line with this notion, pS345 Chk1 levels were reduced in T98G cells treated with 680C91 and there was less robust checkpoint activation in cells co-treated with 680C91 and BCNU. The reduced checkpoint activation was in contrast to the strong induction in pS33 RPA2 signal and increased G2/M arrest observed for cells co-treated with 680C91 and BCNU. These results may be indicative of increased replication-associated DSBs, as phosphorylation of RPA2 by ATR can occur through at least two independent modes – one involving Rad17 and one involving Nbs1 (16). Nbs1-dependent phosphorylation of RPA2 recruits the MRN complex to DNA and involves extensive end-resection. A recent study found that hpol κ protects stalled forks from Mre11 exonuclease activity to promote fork recovery (76). It is possible TDO inhibition in glioma cells leads to down-regulation of hpol κ, which then results in both (a) diminished Chk1 activation and (b) increased resection by the MRN complex. In this regard, the KP could help control the hpol κ-Rad17 arm of the RSR. Disruption of this circuit may lead to a greater dependence on Nbs1-mediated recovery of collapsed forks. A less efficient response to DNA adducts blocking the fork would also explain slowing of the replication machinery and elevated pS33 RPA2 levels in response to BCNU-induced DNA damage. Experiments are ongoing exploring the role of hpol κ in the regulation of replication dynamics in gliomas. While there are many important mechanistic features yet to be discerned, the summation of our results implicates the KP in the resolution of RS inherent to GBM (**Fig. 5G**).

In addition to changes in DNA replication dynamics, we also observed KP-dependent modulation of DNA break repair – with TDO inhibition delaying repair of BCNU-induced strand breaks and the addition of exogenous KYN promoting more rapid clearance of the damage. Recruitment of 53BP1 emerged from our proteomic analysis as a key node in the relationship between TDO activity and DNA repair. This finding was validated by monitoring TDO-dependent changes in 53BP1 foci. The mechanistic features that underlie TDO-dependent changes in 53BP1 dynamics are undoubtedly complex and involve multiple elements (e.g., LCD proteins, trafficking proteins like NUMA1 and NUP153, as well as other proteins directly involved in DNA repair). Of these factors, we explored the notion that impaired SIRT7 activity and localization coincided with defective recruitment of 53BP1 to sites of damage.

SIRT7 participates in multiple aspects of the DDR including chromatin changes and recruitment of DNA repair factors, and loss of SIRT7 sensitizes cells to multiple genotoxic agents (24, 26, 77). SIRT7 is itself recruited to DNA damage sites in a PARP-dependent manner and promotes NHEJ through H3K18 deacetylation and 53BP1 recruitment (61). Furthermore, hyperacetylated p53 accumulates in cells lacking SIRT7, resulting in apoptosis (78). SIRT7 also serves to protect against cell death from persistent DDR by deacetylating ATM late in the DSBR process with failed deacetylation of ATM promoting retention of γH2AX and apoptosis or senescence (23). Based on the results reported here, as well as previously published studies, we propose that activation of the KP in gliomas facilitates SIRT7-mediated recruitment of 53BP1 and subsequent repair of DSBs (**Fig. 5H**). The recruitment of 53BP1 involves multiple players, but it is determined in part by SIRT7-catalyzed deacetylation of H3K18. Blockade of TDO action inhibits this pathway, delaying repair of DSBs and rendering GBM cells more susceptible to BCNU-induced DNA damage. Our study builds upon previous findings to establish a functional link between KP activity in GBM cells and SIRT7-mediated effects on DNA repair – a link that could be related to changes in NAD^+^ supply.

In recent years, the far-reaching impact of NAD^+^-dependent processes on DNA damage, mitochondrial function, neurological disorder, and organismal longevity has received much attention (79–81). Other work has highlighted the role of NAD^+^ levels in resistance to genotoxic anti-cancer drugs. For example, inhibition of the NAD^+^ salvage pathway sensitized the glioblastoma-derived LN428 cell line to TMZ when the nicotinamide phosphoribosyl transferase (NAMPT) inhibitor FK866 was combined with the BER inhibitor methoxyamine (MX) (20). Our results implicate *de novo* NAD^+^ synthesis from tryptophan in a similar phenomenon. Along these same lines, synthesis of NAD^+^ from the tryptophan catabolite QA was also shown to fuel protection against H_2_O_2_, TMZ, and IR (19). The formation of QA depends on expression of quinolinate phosphoribosyltransferase (QPRT), which is normally only expressed in microglial cells. QPRT expression is abnormally high in GBM patients, which may contribute to an increased reliance on *de novo* NAD^+^ supplies for genome protection. Interestingly, T98G cells were the only established cell line shown in a previous report to express QPRT (19), consistent with the augmented effects of BCNU we observed when TDO activity was inhibited.

In conclusion, our finding that inhibition of TDO resulted in failed resolution of RS and delays in repair of BCNU-induced DNA damage carries important implications for how aberrant KP signaling might influence disease progression in gliomas through increased genomic instability and tolerance of therapy-induced DNA damage. Tumors re-wired to express high levels of TDO could have an elevated capacity for tolerating replication stress and DNA damage. The result of this phenomenon might include higher rates of mutagenesis and increased tumor heterogeneity, as well as an enhanced ability to survive genotoxic treatments. Additional work is needed to decipher the exact mechanisms promoting fork recovery and DNA repair in GBM cells with high TDO activity. It will also be interesting to see if similar trends are observed with other clinically relevant genotoxins – including TMZ and IR.

## MATERIALS AND METHODS

### Chemicals

All chemicals were molecular biology grade or better. L-kynurenine (KYN; Cat# K8625) and Carmustine (BCNU; Cat# C0400) were purchased from Sigma-Aldrich (St. Louis, MO). The small molecule inhibitor for TDO (680C91; Cat# 4392) was purchased from Tocris Bioscience (Bristol, UK). For all treatment conditions, final concentration of DMSO/EtOH used was less than 1% (v/v). Experiments were performed with multiple biological replicates, where appropriate experimental treatments were randomized, and all immunofluorescence scoring was conducted in a blinded manner. Statistical evaluations are reported in the figure legends.

### Cell Culture

The glioblastoma-derived cell line T98G was obtained from the American Type Culture Collection (ATCC; Cat# CRL-1690, Manassas, VA). Cells were maintained (5% CO_2_, 37°C) in Modified Eagle’s medium (MEM) containing 10% (v/v) fetal bovine serum and 1% (v/v) antibiotic/antimycotic containing 100 U/mL penicillin, 100 μg/mL streptomycin, and 0.25 μg/mL amphotericin B (Sigma-Aldrich, St. Louis, MO).

### Cell Viability Assay

T98G cell survival was measured using Calcein AM assay, where 5×10^3^ cells were plated per well in a 96-well dish. Cells were treated with 680C91 (10 and 20 μM) for 24 h. Cells were treated for with varying concentrations of BCNU (0 to 2 mM) for an additional 48 h before incubation with Calcein AM dye (2 μM) (Invitrogen; Cat# C1430, Grand Island, NY) at 25°C for 30 m. To determine percent viability, fluorescence values were obtained using Synergy4 plate reader at an excitation wavelength 485 nm and emission wavelength 528 nm. The EC_50_ were calculated using Prism software and relative EC_50_ values were plotted separately.

### Clonogenic Assay

A 6-well dish was plated with 500 cells per well, and cells were allowed to adhere for 24 h. Depending on experimental design, wells were either immediately treated with 10 or 20 μM of 680C91, 60 μM KYN, and 125 μM BCNU for 1 h or pretreated with 680C91/KYN 24 h prior to BCNU treatment. Cells were treated with BCNU (125 μM) for 1 h before replacing with fresh media and allowing cells to recover for 8-10 days. Cells were fixed with formaldehyde (3.7% v/v) before staining with crystal violet (Sigma Aldrich; Cat# V5265, St. Louis, MO). Colonies were counted using an EVOS FL Auto microscope (Life Technologies, Carlsbad, CA) where a colony is considered if at least 25 cells are present. Colony diameter was measured using the microscope described above. For both of these quantifications, three biological replicates were used.

### Flow Cytometry

Cells were stained with Click-it EdU imaging kit (ThermoFisher Scientific; Cat# C10337, Waltham, MA) and the following protocol. Cells were treated with 680C91 (10 or 20 μM) or Kyn (60 μM) 24 h prior to BCNU treatment. Cells were treated with BCNU (125 μM) for 25 h prior to EdU labeling (10 μM) for 1 h, harvested, washed and fixed in 70% (v/v) EtOH before adding permeabilization buffer containing 0.5% (w/v) Triton X-100 in 1x PBS and incubating at RT for 30 min. Cells were washed and incubated in click-it reaction cocktail (click-it reaction buffer, CuSO4, 1x click-it reaction buffer additive, Alexa Fluor 647) for 30 min in the dark at RT. Propidium Iodide (PI) stain (BD Biosciences; Cat# 556463, San Jose, CA) was added for 1 h prior to analysis. During flow analysis, azide only control cells were used for gating. Representative scatterplots for each condition are shown in Figure S1.

### Immunofluorescence

Cells were treated with 680C91 (20 μM) or KYN (60 μM) 24 h prior to BCNU treatment. Cells were treated with BCNU (125 μM) for 24 h before fixing with 3.5% (v/v) formaldehyde and permeabilized with 1x PBS containing 0.2% (w/v) Triton X-100, 0.01% (w/v) sodium azide, 100 μg chicken ovalbumin (Sigma Aldrich; Cat# A2512, St. Louis, MO) and stained for RPA2 (pS33) or γH2AX. Experiments were repeated with in biological triplicate, with at least 130 cells quantified per condition. For the pre-extracted experiments, cells were treated with 0.5% (w/v) Triton-X-100 in 1xPBS on ice for 5 min, prior to fixation with 3.5% (v/v) formaldehyde. Cover slips were permeabilized as described above and costained with 53BP1 and γH2AX. Experiments were repeated in biological duplicate, with at least 159 cells quantified per condition. All experiments were imaged via Olympus Fluoview FV100 microscope and scored via ImageJ and Cell Profiler.

| REAGENT or RESOURCE | DESIGNATION | SOURCE | IDENTIFIER | DILUTION |
| --- | --- | --- | --- | --- |
| **Primary Antibodies** |  |  |  |  |
| Rabbit polyclonal anti-gamma-H2AX (Ser 139) | γ-H2A.X | Abcam | Cat# ab81299 | 1: 1000 |
| Rabbit polyclonal anti-RPA2 (Ser 33) | RPA2(s33) | Novus Biologicals | Cat# NB100-544 | 1:250 |
| Mouse purified anti-H2A.X phospho (Ser139) | γ-H2A.X | BioLegend | Cat# 613401 | 1:200 |
| Rabbit anti-53BP1 | 53BP1 | Cell Signaling Technology | Cat# 4937 | 1:200 |
| **Secondary Antibodies** |  |  |  |  |
| Goat anti-Mouse IgG, DyLight 488 | DyLight 488 | ThermoFisher Scientific | Cat# 35502 | 1:100 |
| Goat anti-Rabbit IgG, Alexa Fluor 647 | AF647 | ThermoFisher Scientific | Cat# A-21244 | 1:100 |

### Immunoblotting

T98G cells were treated with 680C91 or KYN alone, or in combination with BCNU exactly as described earlier, and subsequently processed for immunoblotting. Whole cell lysates (WCL) of T98G cells were prepared by resuspending the frozen cell pellets in 40 mM HEPES (pH 7.5) buffer containing 120 mM NaCl, 1 mM EDT A, 50 mM NaF, and 1% (w/v) Triton X-100. A 1X dilution of protease inhibitor cocktail (Sigma-Aldrich; Cat# P8340, St. Louis, MO) was added to the lysis buffer, as well as the phosphatase inhibitors sodium β-glycerophosphate and sodium orthovanadate. Lysis was achieved by incubation on ice with intermittent mixing for 1 h. This was followed by pelleting the cell debris in a centrifuge by spinning at 21,130xg for 15 min at 4°C. Prior to submission to the UAMS proteomics core facility, harvested cells were separated into nuclear and cytosolic fractions via NE-PER kit following the manufacturer’s instructions (ThermoFisher Scientific; Cat# 78833, Waltham, MA).

To analyze changes in the amount of protein bound to chromatin, we fractionated soluble nuclear and chromatin bound (CB) proteins. Cells were harvested with cell a cell scraper and resuspended in 10 mM HEPES (pH 7.9) buffer containing 10 mM KCl, 1.5 mM MgCl_2_, 0.34 M sucrose, 10% (v/v) glycerol, 0.1% (w/v) Triton X-100, 1 mM DTT, 1 mM NaF, 1 mM Na,VO_4_ and 1X protease inhibitor cocktail. Cells were incubated in lysis buffer for 5 min on ice before centrifugation at 1300xg for 4 min. Nuclei were obtained in the pellet and further separated into soluble and chromatin bound fractions by incubating nuclei in buffer containing 3 mM EDTA, 0.2 mM EGTA, 1 mM DTT, 1 mM NaF, 1 mM Na,VO_4_ and 1X protease inhibitor for 10 min on ice. Supernatant containing soluble nuclear proteins was collected after centrifugation at 1700xg for 4 min. The pellet containing chromatin bound proteins was resuspended in 5x SDS loading dye and boiled for 5 min at ~90°C. To get the chromatin pellet into solution, boiled sample was subjected to water bath sonication until able to easily pipette (~5 min of 30 sec on/ 30 sec off on maximum power).

Total protein concentration of both the WCL and CE samples was estimated by a colorimetric bicinchonic acid (BCA) assay (Thermo Scientific; Cat# 23225, Waltham, MA), using a bovine serum albumin as a standard, as per manufacturer’s instructions.

Proteins from WCL samples were separated on 4-20% (w/v) Tris-Glycine SDS-PAGE gels (Bio-Rad; Cat#456-1094, Hercules, CA). We loaded 30-50 μg protein per sample, and the proteins were separated by electrophoresis under a constant voltage of 120 V for 60-90 m. The gel-separated protein bands were transferred on to a 0.2 μm nitrocellulose (Bio-Rad; Cat# 162-0112, Hercules, CA) or polyvinylidene difluoride (PVDF; Bio-Rad; Cat# 162-0177, Hercules, CA) membrane under constant current of 0.2 A for 60-75 min in an ice-bath. The CE samples were subjected to immunoblotting using the same protocol as described above, when probing for all proteins other than ATM and ATR. The gel-running and transfer protocols were modified for the blots probed for the high molecular weight proteins (>250 kDa) ATM and ATR in the CE samples as follows: the samples were loaded on 4-8% (w/v) Tris-acetate SDS-PAGE gels (ThermoFisher Scientific; Cat# EA03752BOX, Waltham, MA). The running buffer and transfer buffer were prepared as per manufacturer’s instructions. PVDF membrane was used for these blots, and transfer was performed at a constant current of 0.35 A for 4 h at 4°C. After transfer was completed, the blots were incubated in a blocking solution containing 5% (w/v) dried milk solution in 1X Tris-buffered saline (TBS) for 1 h. After blocking, the blots were incubated overnight with the suitable primary antibody on a shaker at 4°C (a complete list of all the primary and secondary antibodies used for immunoblotting is provided in the antibodies resources table). The following day, the blots were washed twice in 1X TBS containing 0.1% (v/v) Tween-20 (TBST), and then incubated in an HRP-conjugated secondary antibody on a shaker at room temperature. After washing twice with TBST, the blots were developed by enhanced chemiluminescence (ECL) using a kit (Bio-Rad; Cat# 1705060, Hercules, CA) and the protein bands were visualized on a ChemiDoc digital imager (Bio-Rad; Cat# 12003153, Hercules, CA). Uncropped immunoblots are shown in **Figures S4** and **S5**.

| REAGENT or RESOURCE | DESIGNATION | SOURCE | IDENTIFIER | DILUTION |
| --- | --- | --- | --- | --- |

| Primary Antibodies |  |  |  |  |
| --- | --- | --- | --- | --- |
| Rabbit polyclonal anti-GAPDH | GAPDH | Cell Signaling Technology | 14C10; Cat# 2118 | 1: 10,000 |
| Rabbit polyclonal anti-Akt | Akt | Cell Signaling Technology | Cat# 9272 | 1: 1000 |
| Rabbit polyclonal anti-phospho-Akt (Ser 473) | p-Akt | Cell Signaling Technology | Cat# 4060 | 1: 1000 |
| Rabbit polyclonal anti-ATM | ATM | Cell Signaling Technology | D2E2; Cat# 2873 | 1: 750 |
| Rabbit polyclonal anti-Ezh2 XP® | Ezh2 | Cell Signaling Technology | D2C9; Cat# 5246 | 1:1000 |
| Rabbit polyclonal anti-histone H3 | H3 | Cell Signaling Technology | D1H2; Cat# 4499 | 1: 2000 |
| Rabbit polyclonal anti-gamma-H2AX (Ser 139) | γ-H2A.X | Abcam | Cat# ab81299 | 1: 1000 |
| Rabbit polyclonal anti-phospho-Chk1 (Ser 345) | p-Chk1 | Cell Signaling Technology | 133D3; Cat# 2348 | 1: 1000 |
| Rabbit polyclonal anti-Chk2 | Chk2 | Abcam | Cat# Ab109413 | 1:1000 |
| Rabbit polyclonal anti-phospho-Chk2 (Thr 68) | p-Chk2 | Cell Signaling Technology | C13C1; Cat# 2197 | 1: 1000 |
| Rabbit polyclonal anti-acetyl histone H3 (Lys 18) | H3 | Cell Signaling Technology | Cat# 9675 | 1: 750 |
| Mouse monoclonal anti-poly (ADP-ribose) polymer | PAR | Abcam | 10H; Cat# ab14459 | 1:1000 |
| Mouse monoclonal anti-PARP-1 | PARP1 | Santa Cruz Biotech | F2; Cat# sc-8007 | 1:1000 |
| Mouse monoclonal anti-SIRT-7 | SIRT7 | Santa Cruz Biotech | C3; Cat# sc-365344 | 1: 1000 |
| Rabbit anti-Acetylated-Lysine | panAc-Lys | Cell Signaling Technology | Cat# 9441S | 1:1000 |
| Secondary Antibodies |  |  |  |  |
| Goat polyclonal anti-Rabbit HRP conjugated | Anti-rabbit | ThermoFisher Scientific | Cat# 32460 | 1:2000 – 1:5000 |
| Goat polyclonal anti-Mouse HRP conjugated | Anti-mouse | ThermoFisher Scientific | Cat# 32430 | 1:2000 – 1:5000 |

### Micronucleation

Micronucleation experiments were performed as previously described (13). Briefly, cells were treated with 680C91 (20 μM) or KYN (60 μM) 24 h prior to BCNU treatment. Cells were treated with BCNU (125 μM) for 24 h before treating with cytochalasin-B (2 μg/mL) (Sigma Aldrich; Cat# 250233, St. Louis, MO) for an additional 24 h. Slides were stained with α-tubulin. Images were obtained via Olympus Fluoview FV100 microscope and subjected to blinded scoring in biological triplicate.

| REAGENT or RESOURCE | DESIGNATION | SOURCE | IDENTIFIER | DILUTION |
| --- | --- | --- | --- | --- |
| **Primary Antibodies** |  |  |  |  |
| Mouse monoclonal anti-α-Tubulin | α-Tubulin | MiliporeSigma | Cat# T9026 | 1:500 |
| **Secondary Antibodies** |  |  |  |  |
| Goat anti-Mouse IgG, DyLight 650 | DyLight 650 | ThermoFisher Scientific | Cat# 84545 | 1:50 |

### DNA fiber spreading

Cells were pre-treated with 680C91 (20 *μ*M) or KYN (60 *μ*M) 24 h prior to start of experiment. Pre-treated cells were incubated with media containing 20 *μ*M 5-chloro-2’-deoxyuridine (CldU) (Milipore Sigma; Cat# C6891, St. Louis, MO) for 30 m at 37°C before being washed and incubated in media containing 100 *μ*M 5-iodo-2’-deoxyuridine (IdU) (Milipore Sigma; Cat# I7125, St. Louis, MO) alone or in combination with BCNU (125 *μ*M) for 1 h at 37°C. Fork restart experiments were also pretreated as described above followed by 3mM hydroxyurea (HU) (Milipore Sigma; Cat# H8627, St. Louis, MO) treatment for 4 h after CldU treatment and then 30 min IdU pulse. Cells were counted and diluted to 1 million cells/mL and mixed 1:1 with unlabeled cells. Around 2 μL of cell suspension was dropped onto top of glass slides and solution was left to air dry until volume was slightly reduced before mixing in DNA lysis buffer (0.5% (w/v) SDS, 200 mM Tris-HCl pH 7.4, 50 mM EDTA) and incubating for 5 min at RT. Slides were then tilted to ~15° for the solution to reach to bottom of the slides and then dried. Dried slides were fixed in 3:1 dilution of methanol and acetic acid for 3 min before incubating in 2.5 M HCl for 70 min. Slides were washed and transferred into a humid box where they were incubated in blocking buffer (10% goat serum in 1x PBS) for 60 min at RT. Slides were stained for CldU and IdU for 2 h at RT before adding corresponding secondary antibodies for an additional h at RT. The length of IdU and CldU was measured for each fiber counted and normalized to analogue incubation time (60 min or 30 min respectively). Images were obtained via Nikon Ti2 Eclipse microscope and subjected to blinded scoring in biological triplicate. Statistical analysis performed was a non-parametric Mann-Whitney analysis of means.

| REAGENT or RESOURCE | DESIGNATION | SOURCE | IDENTIFIER | DILUTION |
| --- | --- | --- | --- | --- |
| **Primary Antibodies** |  |  |  |  |
| Rat anti-Bromodeoxyuridine | anti-BrdU (CldU) | Novus Biologicals | Cat# NB500-169 | 1:250 |
| Mouse anti-Bromodeoxyuridine | anti-BrdU (IdU) | BD Biosciences | (B44) Cat# 347580 | 1:250 |
| **Secondary Antibodies** |  |  |  |  |
| Goat anti-Mouse IgG, DyLight 488 | DyLight 488 | ThermoFisher Scientific | Cat# 35502 | 1:100 |
| Goat anti-Rat IgG, Alexa Fluor 594 | AF594 | Invitrogen | Cat# A-11007 | 1:100 |

### FASP bHPLC TMTMass Spectrometry

T98G cells (~10^6^ cells/condition) were pre-treated with 680C91 (10 or 20 μM) or simply cultured in media for 24 h prior to BCNU treatment. After the 680C91 pretreatment, the media containing the TDO inhibitor was replaced with either the standard MEM/FBS/anti-anti media or with media containing BCNU (125 μM). After 24 h treatment with either BCNU or untreated media, the cells were harvested and subjected to nuclear and cytosolic fractionation via NE-PER kit (ThermoFisher Scientific; Cat# 78833, Waltham, MA). Nuclear protein was isolated and 100 μg of lysate was submitted to UAMS proteomics core facility.

Purified proteins were reduced, alkylated, and digested using filter-aided sample preparation (82). Tryptic peptides were labeled using a tandem mass tag 10-plex isobaric label reagent set (Thermo) following the manufacturer’s instructions. Labeled peptides were separated into 36 fractions on a 100 x 1.0 mm Acquity BEH C18 column (Waters) using an UltiMate 3000 UHPLC system (Thermo) with a 40 min gradient from 99:1 to 60:40 buffer A:B ratio under basic pH conditions, and then consolidated into 12 superfractions. Each super-fraction was then further separated by reverse phase XSelect CSH C18 2.5 um resin (Waters) on an in-line 120 x 0.075 mm column using an UltiMate 3000 RSLCnano system (Thermo). Peptides were eluted using a 60 min gradient from 98:2 to 67:33 buffer A:B ratio. Eluted peptides were ionized by electrospray (2.25 kV) followed by mass spectrometric analysis on an Orbitrap Fusion Lumos mass spectrometer (Thermo) using multi-notch MS3 parameters. MS data were acquired using the FTMS analyzer in top-speed profile mode at a resolution of 120,000 over a range of 375 to 1500 m/z. Following CID activation with normalized collision energy of 35.0, MS/MS data were acquired using the ion trap analyzer in centroid mode and normal mass range. Using synchronous precursor selection, up to 10 MS/MS precursors were selected for HCD activation with normalized collision energy of 65.0, followed by acquisition of MS3 reporter ion data using the FTMS analyzer in profile mode at a resolution of 50,000 over a range of 100-500 m/z.

Proteins were identified and TMT MS3 reporter ions quantified by searching the *Homo sapiens* database using MaxQuant (version 1.6.2.10, Max Planck Institute) with a parent ion tolerance of 3 ppm, a fragment ion tolerance of 0.5 Da, and a reporter ion tolerance of 0.001 Da. Protein identifications were accepted if they could be established with less than 1.0% false discovery and contained at least 2 identified peptides. Protein probabilities were assigned by the Protein Prophet algorithm (83).

TMT MS3 reporter ion intensity values were log_2_ transformed and missing values were imputed by a normal distribution for each sample using Perseus (Max Planck Institute). TMT batch effects were removed using ComBat (84) in order to correct for technical variation due to multiplexing samples across multiple TMT10plex batches. Statistical analysis was performed using Linear Models for Microarray Data (limma) with empirical Bayes (eBayes) smoothing to the standard errors (85). Proteins with an FDR adjusted p-value < 0.05 and a fold change > 2 were considered to be significant. Significant proteins were analyzed for protein networks and pathways using the Ensemble of Gene Set Enrichment Analyses (EGSEA) Bioconductor package (86) and Qiagen’s Ingenuity Pathway Analysis (87).

### Alkaline Comet Assay

The assay was conducted using the Comet Assay kit (Trevigen; Cat# 4252-040-K, Gaithersburg, MD). Briefly, 100,000 cells per condition were plated in a 12 well dish and pre-treated for 24 h with either 680C91 (20 μM) or KYN (60 μM). Cells were treated with BCNU (125 μM) for 24 h before being harvested or let to recover for an additional 24 h ± 680C91/KYN. Harvested cells were washed in ice cold 1x PBS, and counted. Cells were diluted to 100,000 cells/mL and resuspended in LMAgarose to a final plated dilution of around 200-300 cells per well on the comet slide. The comet slide with wells coated with cell/agar mixture was incubated at 4°C in the dark to allow for the agar to solidify before immersing slide in proprietary lysis solution at 4 °C for 1 h. The comet slide was immersed in alkaline unwinding solution (200 mM NaOH, 1 mM EDTA in deionized H_2_O) for 20 min at RT before placing in the electrophoresis unit in ~850 mL of alkaline electrophoresis solution (200 mM NaOH, 1 mM EDTA in deionized H_2_O) and running at 21 V for 30 min at 4°C. The comet slide was washed and dried completely at 37°C before staining with SYBR gold (Sigma Aldrich; Cat# S9430, St. Louis, MO) diluted in TE buffer (10 mM Tris-HCl pH 7.5, 1 mM EDTA). Comets were visualized using the EVOS microscope and scored using CometScore software. At least 190 comets were scored from at least two biological replicates, and tail moment was analyzed using one-way ANOVA with Tukey post-test.

### Data Availability

All source data are stored on a secure UAMS server and can be made available upon request. Proteomics results are available in the Proteomics Identification Database (PRIDE) repository (https://www.ebi.as.uk/pride/).

## Supporting information

Supplemental Material for Reed et al

## Acknowledgements

We are grateful to Ms. Andrea Harris in the UAMS Flow Cytometry Core and Mr. Brian Koss for assistance with the analysis and presentation of the flow cytometry results. This work was supported in part by National Institutes of Health (NIH) grants CA183895 (R.L.E.) and GM121293 (A.J.T.) with additional support from the Arkansas Breast Cancer Research Program, a Seeds of Science award from the Little Rock Envoys, a Barton Bridging Award from the UAMS College of Medicine, and the University of Arkansas for Medical Sciences Translational Research Institute grant (UL1 TR003107) through the National Center for Advancing Translational Sciences of the NIH. The content is solely the responsibility of the authors and does not necessarily represent the official views of the NIH.

## Author contributions

RLE, MRR, and SDB designed the research. MRR, LM, AK, MKZ, and ACLB performed the experiments. MRR, MKZ, LM, AK, SDB and RLE analyzed the experimental results. MRR, AK, SDB, and RLE wrote the paper. LM, MKZ, ACLB, and AJT provided comments on the manuscript. All authors approved the final version of the manuscript.

