## Supplemental Material for Reed et al for "Inhibition of tryptophan 2,3-dioxygenase impairs DNA damage tolerance and repair in glioma cells"

*Running Title: Kynurenine signaling and genome maintenance in gliomas*

**TABLE OF CONTENTS**

**TABLE S1. List of proteins identified by TMT mass spectrometry across all samples.**

**TABLE S2. IPA comparing proteomic results for T98G cells treated with either DMSO or 10 μM 680C91.**

**TABLE S3. IPA comparing proteomic results for T98G cells treated with either DMSO or 20 μM 680C91.**

**TABLE S4. IPA comparing proteomic results for T98G cells treated with either DMSO or 125 μM BCNU.**

**TABLE S5. IPA comparing proteomic results for T98G cells treated with either 125 μM BCNU or 20 μM 680C91.**

**TABLE S6. IPA comparing proteomic results for T98G cells treated with either DMSO or 125 μM BCNU and 20 μM 680C91.**

**TABLE S7. IPA comparing proteomic results for T98G cells treated with either 125 μM BCNU alone or 125 μM BCNU and 20 μM 680C91.**

**FIGURE S1. Representative flow cytometry scatterplots for cells treated with either 680C91 (10 or 20 μM), KYN (60 μM), BCNU (125 μM) or combination treatments.**

**FIGURE S2. Quantification of changes in protein abundance as a function of molecular weight.**

**FIGURE S3. Immunoblot analysis of acetylated lysine, PAR-ylation, and AKT activation in response to KP modulation.**

**FIGURE S4. Uncropped immunoblots for whole cell lysate (WCL) samples.**

**FIGURE S5. Uncropped immunoblots for chromatin-bound (CB) samples.**

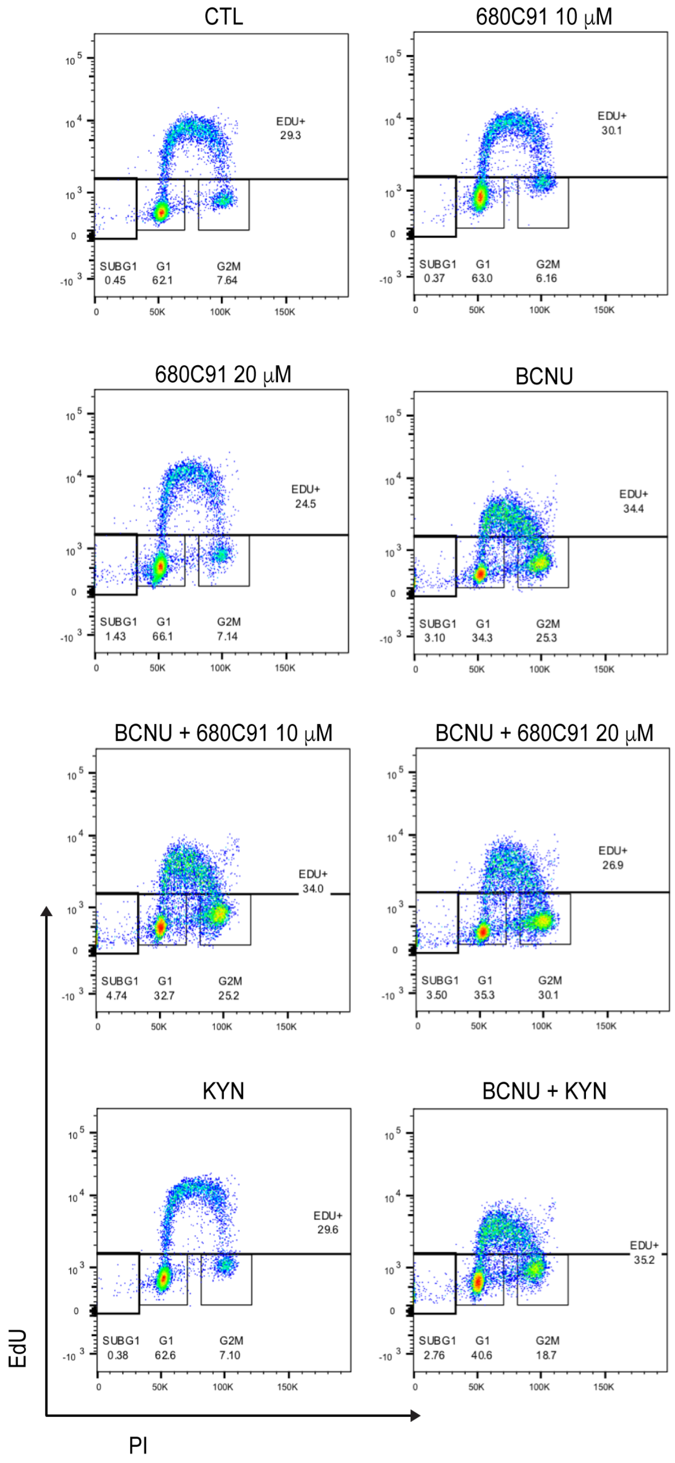


**FIGURE S1. Representative flow cytometry scatterplots for cells treated with either 680C91 (10 or 20 μM), KYN (60 μM), BCNU (125 μM), or combination treatments.** Representative scatterplots and gating profiles are shown for experiments that are summarized in Figure 3B of the main text.

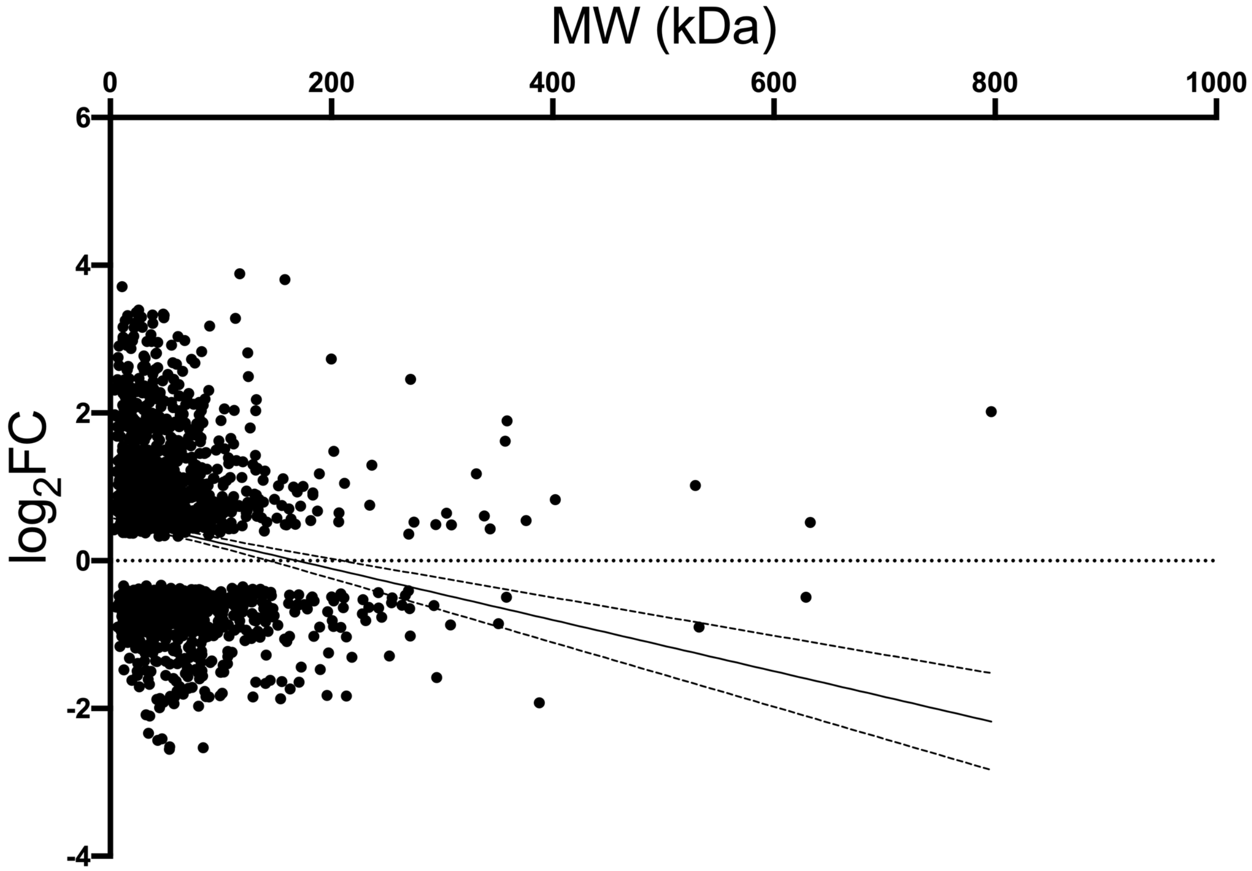


**FIGURE S2. Quantification of changes in protein abundance as a function of molecular weight.** The $log_2$ fold-change (FC) for all proteins with an FDR-adjusted p-value of <0.05 are shown plotted as a function of molecular weight (MW) in kDa. Simple linear regression was used to provide an initial directionality to the resulting plot. The resulting line is shown along with the 95% confidence intervals (dashed lines). To evaluate correlation between the direction of change and protein MW, a Pearson correlation value was calculated using GraphPad Prism, resulting in a Pearson r value of -0.1852 (95% confidence interval of -0.2316 to -0.1380, $R^2 = 0.034$) and a two-tailed P value of <0.0001.

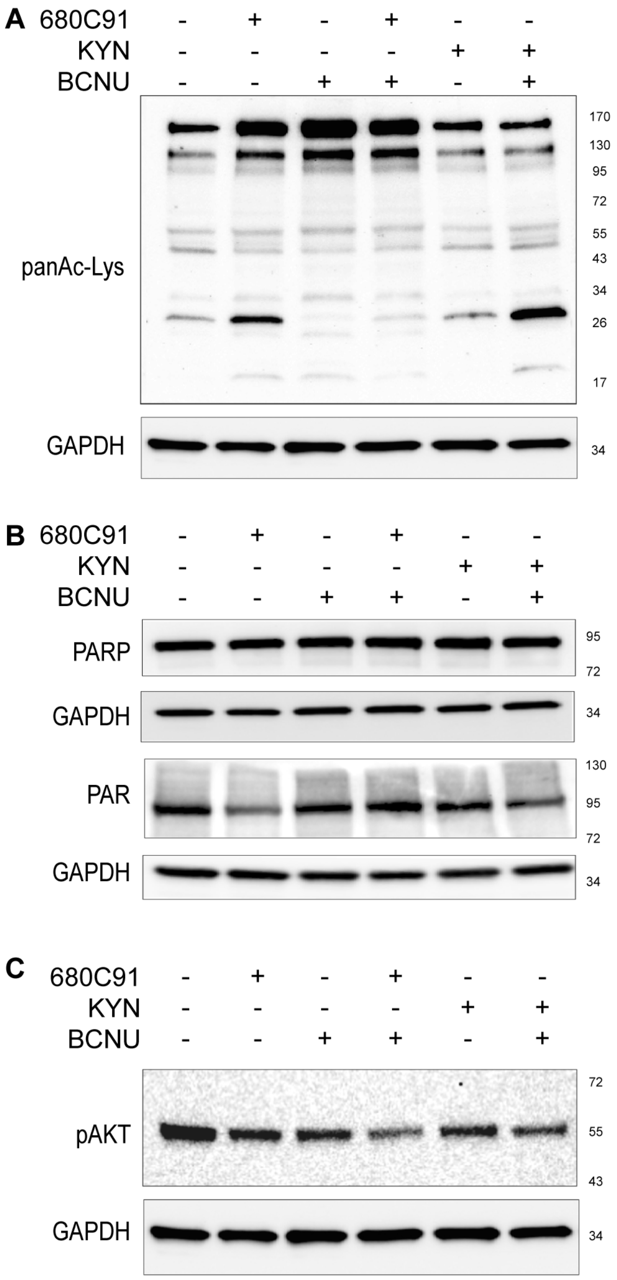


**FIGURE S3. Immunoblot analysis of acetylated lysine, PAR-ylation, and AKT activation in response to KP modulation.** Immunoblot of (A) panAcetyl-Lysine expression, (B) total PARP and PAR-ylation levels, and (C) phosphorylated AKT (S473).

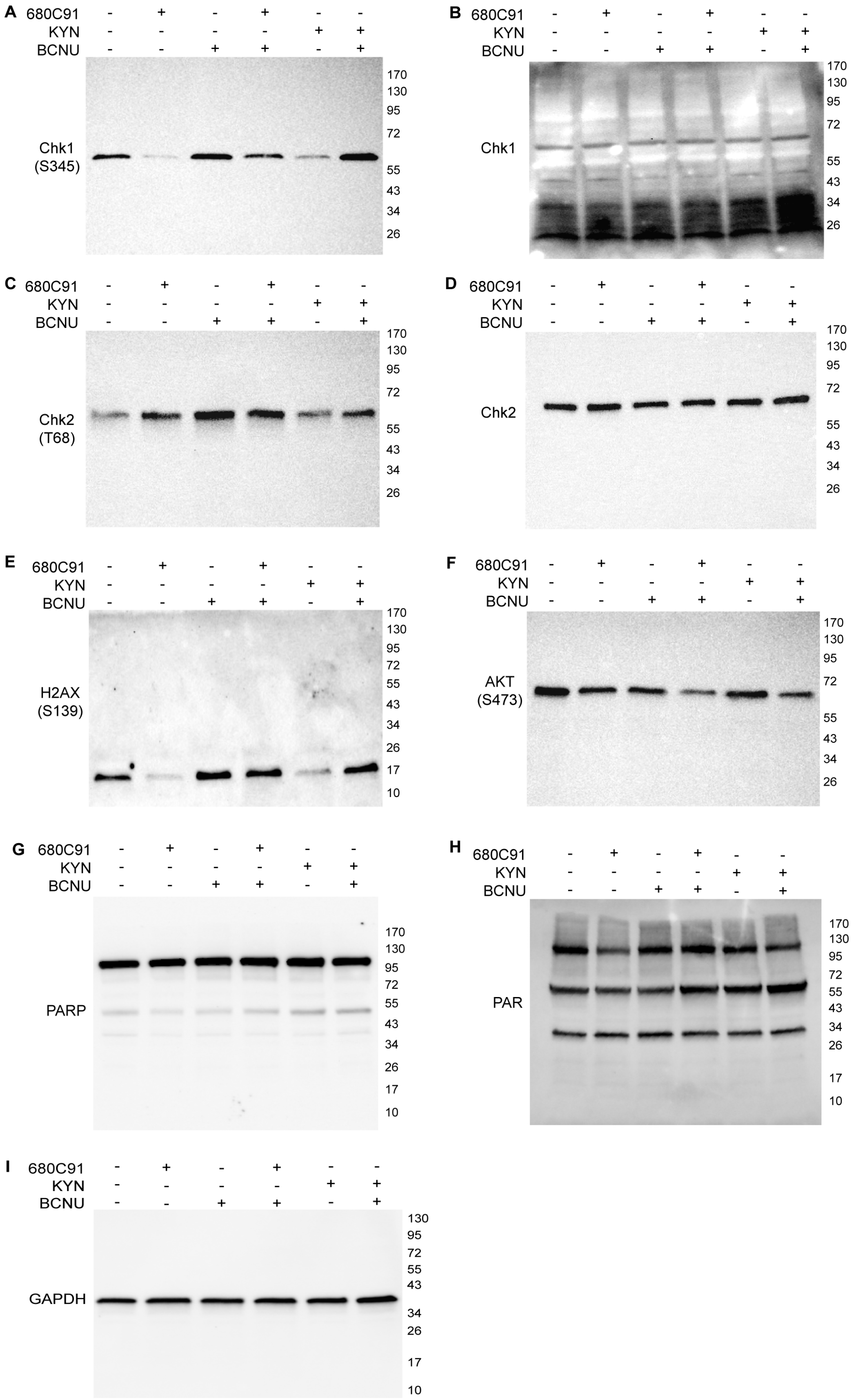
A
680C91 - + - + - -
KYN - - - - + +
BCNU - - + + - +
Chk1 (S345)
170 130 95 72 55 43 34 26
B
680C91 - + - + - -
KYN - - - - + +
BCNU - - + + - +
Chk1
170 130 95 72 55 43 34 26
C
680C91 - + - + - -
KYN - - - - + +
BCNU - - + + - +
Chk2 (T68)
170 130 95 72 55 43 34 26
D
680C91 - + - + - -
KYN - - - - + +
BCNU - - + + - +
Chk2
170 130 95 72 55 43 34 26
E
680C91 - + - + - -
KYN - - - - + +
BCNU - - + + - +
H2AX (S139)
170 130 95 72 55 43 34 26 17 10
F
680C91 - + - + - -
KYN - - - - + +
BCNU - - + + - +
AKT (S473)
170 130 95 72 55 43 34 26
G
680C91 - + - + - -
KYN - - - - + +
BCNU - - + + - +
PARP
170 130 95 72 55 43 34 26 17 10
H
680C91 - + - + - -
KYN - - - - + +
BCNU - - + + - +
PAR
170 130 95 72 55 43 34 26 17 10
I
680C91 - + - + - -
KYN - - - - + +
BCNU - - + + - +
GAPDH
130 95 72 55 43 34 26 17 10

**FIGURE S4. Uncropped immunoblots for whole cell lysate (WCL) samples.** Images of full blots of the samples shown as cropped images in Figure 1H and Figure S3. The images shown are one of two replicates. The primary antibody used for probing each blot is indicated to the left of each panel, and the positions of bands for the reference molecular weight ladder are indicated towards the right of each blot.

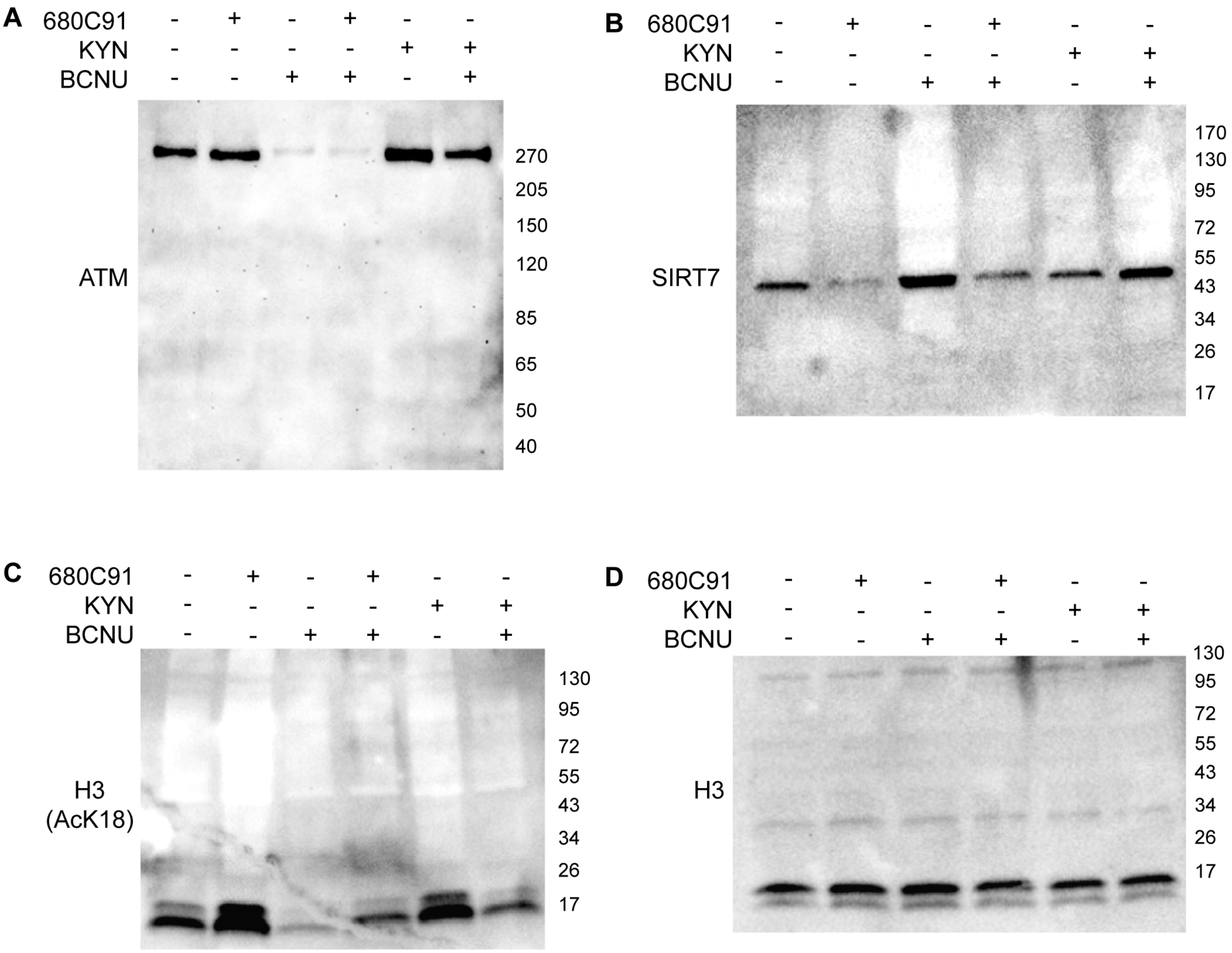


**FIGURE S5. Uncropped immunoblots for chromatin-bound (CB) samples.** Images of full blots of the samples shown as cropped images in Figure 5 of the main text. The primary antibody used for probing each blot is indicated above each panel, and the positions of bands for the reference molecular weight ladder are indicated towards the left of each blot.
